# The effect of *Sinorhizobium meliloti* volatilomes and synthetic long-chain methylketones on soil and *Medicago truncatula* microbiomes

**DOI:** 10.1101/2025.10.01.679752

**Authors:** Pieter van Dillewijn, Lydia M. Bernabéu-Roda, Virginia Cuéllar, Rafael Núñez, Otto Geiger, Isabel M. López-Lara, María J. Soto

**Affiliations:** Department of Biotechnology and Environmental Protection, Estación Experimental del Zaidín, CSIC, Granada, Spain; Scientific Instrumentation Service, Estación Experimental del Zaidín, CSIC, Granada, Spain; Centro de Ciencias Genómicas, Universidad Nacional Autónoma de México, Cuernavaca, Mexico

**Keywords:** Bacterial volatile compounds, methylketones, volatilome, microbiome, soil-, rhizosphere-, root endosphere-bacterial communities, *Ensifer*/*Sinorhizobium*, *Medicago truncatula*

## Abstract

Bacterial volatile compounds play important roles in intra- and interkingdom interactions but very little is known about their effects on soil and plant microbiomes. The legume symbiont *Sinorhizobium meliloti* (Sm) releases volatile methylketones (MKs), one of which acts as an infochemical in bacteria and hampers plant-bacteria interactions. MK production in Sm is modestly increased in the absence of the long-chain-fatty-acyl-coenzyme-A (CoA) synthetase FadD. To explore further the ecological role of MKs on soil and plant bacterial communities, we aimed at obtaining an MK-overproducer Sm strain by deleting the 3-oxo-acyl-CoA-thiolase-encoding *fadA* gene. Analyses of the Sm wild type (wt), and *fad* mutant volatilomes identified seventeen compounds consisting mostly of MKs and fatty acid methyl esters (FAMEs) and revealed that the *fadA* mutant produced more MKs than the *fadD* mutant and much more than the wt, while in the *fadD* mutant FAME emission was increased. When natural soil or the rhizosphere of *Medicago truncatula* were exposed to wt and *fadA* volatilomes or synthetic MKs, bacterial alpha- or beta-diversity were not strongly affected but specific genera were identified which responded differentially to each condition. Interestingly, Sm volatilomes had a significant effect on root endosphere *Ensifer*/*Sinorhizobium* populations by maintaining their abundance over time in contrast to control conditions or exposure to synthetic MKs. This study provides new insights on the synthesis of rhizobial volatile compounds and represents the first exploration of the effects of bacterial volatilomes on plant bacterial communities, contributing to increase our knowledge on the complex molecular bases underlying plant-bacteria interactions.

## Introduction

Bacteria release a large number of low molecular weight volatile compounds (VCs) including inorganic volatile compounds (VICs) such as CO_2_, NO, H_2_, H_2_S, NH_3_, or HCN and organic volatile compounds (VOCs) belonging to various chemical classes, such as terpenes, nitrogen- and sulfur-containing compounds, hydrocarbons, ketones, alcohols, aldehydes, or acids (Audrain et al. 2015; Kemmler et al. 2025). Bacterial VCs are well known for their beneficial effects on plants where they can increase plant growth and/or resistance to biotic and abiotic stresses but they can also play important roles in microbe-microbe interactions by affecting growth and bacterial behaviors such as motility, biofilm formation, virulence, or stress and antibiotic resistance (Schulz-Bohm et al. 2017; Sharifi et al. 2022; Weisskopf et al. 2021). However, very little is known of how discrete volatiles or bacterial volatile blends, also known as volatilomes, affect soil and plant associated microbiomes, which could have a great influence on plant health. The existing limited research has focused mostly on the effect of exogenous applied gases on soil or plant microbiomes but usually at elevated levels, for instance within the context of global climate change rather than as bacterially produced VICs (He et al. 2020; Yu and Chen 2019; Wang et al. 2020). With respect to bacterial emitted VOCs, we are not aware of any study focused on the effect of discrete bacterial emitted VOCs as airborne transmitted compounds on soil or plant microbiomes. The closest is a study in which *N*,*N*- dimethylhexadecylamine (DMHDA), a VOC produced by some beneficial rhizobacteria, was applied as solution into the growth medium of *in vitro* grown *Medicago truncatula* which had been inoculated with the microbial fraction from soil (Real-Sosa et al. 2022). In that study it was found that the presence of DMHDA modulated the shoot and root endophytic bacterial communities. With respect to bacterial volatilomes or volatile blends, to date and to the best of our knowledge, the only study found is that of the effect of the volatilome emitted by *Bacillus amyloliquefaciens* NJN-6 on a soil microbiome which was shown to alter soil microbial communities (Yuan et al. 2017). However, studies of the effect of bacterial volatilomes on plant associated microbiota such as those of the rhizosphere or endosphere are not available.

Soil-dwelling bacteria collectively referred to as rhizobia are able to colonize and invade legume roots, leading to the formation of a specialized organ, the nodule, inside of which the bacteria reduce nitrogen to ammonia that can be used by the plant (Poole et al. 2018). Despite being one of the best-known plant-interacting microorganisms and their relevance for their ability to establish beneficial nitrogen fixing symbiosis with legumes, the knowledge about the production and ecological role of VCs emitted by rhizobia is limited (Soto et al. 2021).

The legume symbiont *Sinorhizobium meliloti* (Sm) GR4, which forms nitrogen fixing nodules on *Medicago* plants such as alfalfa (*Medicago sativa*) or barrel medic (*Medicago truncatula*) was shown to emit the volatile long-chain methylketones (MKs): 2- tridecanone (2-TDC), 2-pentadecanone (2-PDC), and 2-heptadecanone (2-HpDC) (López-Lara et al. 2018). Two of these MKs were shown to affect bacterial surface motility when applied as volatile pure chemicals and one of them, 2-TDC, was shown to be an infochemical that affects important bacterial traits and hampers plant–bacteria interactions by interfering with microbial colonization of plant tissues not only of Sm but also of other bacterial species (López-Lara et al. 2018). However, the effect of Sm emitted MKs, separately or in volatilomic blends on soil and plant associated micobiomes remains unknown.

In Sm, the inactivation of the long-chain fatty acyl-Coenzyme A ligase FadD, leads to a modest accumulation of MKs (López-Lara et al. 2018). Although information on how bacteria produce MKs is scarce, most likely, MKs produced by *fadD* mutants derive from substrates and enzymatic reactions similar to those described for tomato plants (Yu et al. 2010). Namely, the fatty acid biosynthesis intermediates, 3-oxo-acyl-acyl carrier proteins (3-oxo-acyl-ACPs), might be hydrolyzed by dedicated thioesterases resulting in the release of 3-oxo fatty acids, which enzymatically or spontaneously decarboxylate to generate the corresponding MK products (Fig. S1). Recently, the YbgC-like thioesterase SMc03960 has been shown to contribute to MK production in the alfalfa symbiont (Bernabéu-Roda et al. 2025).

MKs may also derive from 3-oxo-acyl-CoAs, which are intermediates of the fatty acid degradation pathway (Fig. S1). Indeed, overproduction of MKs has been achieved in bacteria by modifying the β-oxidation cycle to accumulate 3-oxo-acyl-CoAs, together with the overexpression of a thioesterase (Goh et al. 2012, 2014). Based on these studies, and in order to obtain an Sm strain that overproduces MKs, the *fadA* gene coding for a 3- oxo-acyl-CoA thiolase was deleted in Sm GR4. The VOC compositions of the volatilomes emitted by Sm GR4 (wt) and its *fadD* and *fadA* derivative mutants have been determined. Since the aim of this study was to elucidate the effect of MKs on natural bacterial populations, Sm wild type and MK-enriched volatilomes, as well as synthetic MKs, were tested for their effects on soil and plant-associated microbiomes for the first time.

## Materials and Methods

### Chemicals, bacterial strains, plasmids, and growth conditions

All reagents and chemicals including 2-tridecanone (2-TDC), 2-pentadecanone (2- PDC) and 2-heptadecanone (2-HpDC), were obtained from Sigma-Aldrich (Steinheim, Germany).

The *S. meliloti* (Sm) wild type strain GR4 (Casadesús and Olivares 1979), and its *fadD* and *fadA* derivative mutants were used in this study. The *fadD* mutant strain, GR4FDCSS, was described previously (Amaya-Gómez et al. 2015). The *fadA* mutant strain, GR4FADA, was obtained by replacing the wild type *fadA* (*smc02228*) allele with an unmarked in-frame deleted version, which was generated *in vitro* by overlap extension PCR (Ho et al. 1989) using primers fadA1 (5’- CG<u>GGATCC</u>GCCTATCAGAATGCTGCG-3’, underlined *Bam*HI recognition sequence), fadA2 (5’-CAGCTCGTCGAGCACGTCCTTCTTGCCACG-3’), fadA3 (5’- CGTGGCAAGAAGGACGTGCTCGACGAGCTG-3’), and fadA4 (5’-AAA<u>GAATTC</u>GAGTTCCTGCATCGAGAC-3’, underlined *Eco*RI recognition sequence), and the proofreading Phusion high-fidelity DNA polymerase (Thermo Scientific, Waltham, Massachusetts, USA). The resulting fusion product, in which a deletion of 1050 bp was generated in the *fadA* coding sequence, was cloned into pCR2.1- TOPO (Invitrogen), sequenced, and then subcloned into the suicide vector pK18*mobsacB* (Schäfer et al. 1994) as a *Bam*HI-*Eco*RI fragment. The resulting construct pK18-ΔfadA was introduced into Sm GR4 via biparental mating using *Escherichia coli* S17-1 (Simon et al. 1983), and allele replacement events were selected as described previously (Schäfer et al. 1994). Putative deletion mutant strains were identified by PCR and checked by Southern hybridization using a *fadA*-specific probe.

*E. coli* strains were grown in lysogeny broth (LB) medium (Sambrook et al. 1989) at 37°C. Sm strains were grown at 30 °C in tryptone-yeast extract (TY) medium (Beringer 1974) or in minimal medium (MM) (Robertsen et al. 1981). For solid MM, 1% Difco® Noble Agar (BD, Le Pont de Claix, France) was used. When required the following antibiotics were added (final concentrations): kanamycin (50 μg. mL^-1^ for *E. coli*), streptomycin (50 μg mL^-1^ for *E. coli*, 200 μg mL^-1^ for *S. meliloti*), or spectinomycin (100 μg. mL^-1^ for *S. meliloti*).

### Analyses of volatiles by headspace solid-phase microextraction-GC/MS (HS-SPME-GC/MS)

Organic volatile compounds produced by Sm strains were analyzed as described by (López-Lara et al. 2018). In short, cultures grown in TY broth to the late exponential phase were washed, concentrated and then inoculated onto the surface of 3 ml of MM (1% agar) in 10 ml glass vials equipped with silicone septa and incubated at 30°C for 2 days. Volatiles were collected from the headspace of the vials by SPME with a Divinylbenzene/Carboxen/Polydimethylsiloxane (DVB/CAR/PDMS) coated StableFlex SPME 50/30 mm fiber (Supelco 57298-U) introduced into the headspace of the vial and incubated at 30°C for 60 min. Then the fiber was inserted into the injector of a gas chromatograph/mass spectrometer (GC/MS) (Varian 450GC 240MS, Ion Trap) and separated in a DB5MS-UI column (30 m, 0.25-mm inside diameter, 0.25 µm) under the conditions described by López-Lara et al. (2018). Two methods were used for compound identification. Comparison of mass spectra of detected peaks with those in NIST 17 library and comparison of experimental and theoretical Retention Indexes (*n*-alkane scale). VOC abundance was determined using arbitrary counts after subtraction of compounds detected in the vials containing only medium without bacterial inoculation (blank control).

### Microcosm experiments and sampling

Microcosm experiments were performed as described in Section S1 and Fig. S2. For soil experiments, three replicate pots were taken initially (T=0) or after 1 and 7 days of incubation. To obtain soil samples, the contents of each pot were thoroughly mixed, placed in 50 mL tubes and stored at -20°C until DNA extraction. For plant assays, three replicate planted pots were taken initially (T=0) or after 14 days of incubation and rhizosphere and root endosphere samples were obtained as described in Section S2.

### Total DNA isolation and amplicon sequencing analysis

DNA from soil, rhizosphere and root endosphere (root powder) was obtained from 3 replicate pots using the Fast DNA Spin Kit for Soil (MP Biomedicals LLC, Solon, Ohio, USA). Purity and DNA concentrations were determined using Nanodrop ND-1000 (Thermo Fisher Scientific) and Qubit dsDNA BR Assay kit (Live Technologies, Invitrogen, USA). Paired end amplicon sequencing (2 x 275 bp) of the V3-V4 region of the 16S ribosomal RNA gene was performed using MiSeq Illumina technology. Prior to library preparation of endosphere samples, treatment with PCR clamps was performed to reduce plant chloroplast and mitochondria 16S rRNA amplification (Lundberg et al. 2013). The sequencing data have been deposited in the NCBI Sequence Read Archive (SRA) under BioProject accession ID PRJNA1336214.

### Bioinformatic data analysis

16S rRNA amplicon sequences were processed using QIIME2 version 2024.5 (https://qiime2.org)) (Bolyen et al. 2019) as described in van Dillewijn et al. (2025) and detailed in Section S3 to obtain features or amplified sequence variants (ASVs). In order to detect differentially abundant genera between the different treatments, soil types and time points, ANCOM-BC (Lin and Peddada 2020) with an adjusted p-value (q-value) < 0.05 was used within the QIIME2 package. One-way analysis of variance (ANOVA) at a significance level of 0.05 and Welch t-test were calculated using the Excel program from Microsoft Office (2019) (Microsoft).

## Results

### Volatilome profile of *S. meliloti* GR4 (wt) and its *fadD* and *fadA* mutant derivatives

As previously mentioned in the Introduction, bacterial long-chain methylketones (MKs) may derive from intermediates of the fatty acid biosynthesis or degradation pathways, which makes it difficult to obtain an MK-null mutant. To evaluate the effect of MKs present in the volatilome of *S. meliloti* on soil and plant bacterial populations, an MK-overproducer could be used as an alternative. Inactivation of *fadD* in Sm leads to a modest increase in MK production (López-Lara et al. 2018). Therefore, we aimed at obtaining a strain exhibiting higher release of volatile MKs by generating a *fadA* mutant. The working hypothesis was that the activity of endogenous thioesterases on accumulated 3-oxo-acyl-CoAs in this strain would generate the corresponding 3-oxo-acids that, by spontaneous or enzymatic decarboxylation, would release MKs (Fig. S1).

To determine to what extent the *fadA* mutant overproduces MKs, VCs emitted by cells growing on solid MM were determined by headspace SPME-GC/MS and compared to the volatilome profile emitted by the wild type strain (wt) and the *fadD* mutant under the same growth conditions. As shown in Fig. 1, 17 VOCs were detected including 2 alcohols, 6 MKs (Fig. 1a) and 9 fatty acid methyl esters (FAMEs; Fig. 1b). Significantly larger (p<0.05, Welch t- test) amounts of 2-tridecanol (more than 7 fold), 2-TDC (more than 10-fold), 2-PDC and 2-HpDC (more than 30-fold each) and octadecenoic acid methyl ester (almost 2-fold) were detected in the volatilome produced by the *fadA* mutant than in that of the wild type. As reported previously (López-Lara et al. 2018), compared to the wild type strain, the *fadD* mutant exhibits increased production (approx. 4-fold more) of the MKs 2-TDC, 2-PDC and 2-HpDC, but their levels were well below those emitted by the *fadA* mutant (Fig. 1a). Compared to the wild type strain, the *fadD* mutant also emits more 2-tridecanol (2-fold more) but especially significantly larger amounts of the following FAMEs: decanoic acid methyl ester (almost 5-fold increase), dodecanoic acid methyl ester (about 2-fold increase), tetradecanoic acid methyl ester (about 2-fold increase), hexadecanoic acid methyl ester (almost 3-fold increase), and octadecenoic acid methyl ester (almost 4-fold increase), as well as large but not significant increases of octanoic acid methyl ester (about 2-fold increase), and nonanoic acid methyl ester (about 17-fold increase) (Fig. 1b). These results indicate that, indeed, the mutation of *fadA* leads to MK overproduction but does not increase much the emission of FAMEs while the mutation of *fadD* increases modestly the release of MKs but much more the emission of FAMEs. Since this study focused on the effect of MKs on soil and plant associated microbiomes, further studies were performed with the wild type and the MK- overproducing *fadA* mutant strain.

**Fig. 1.**
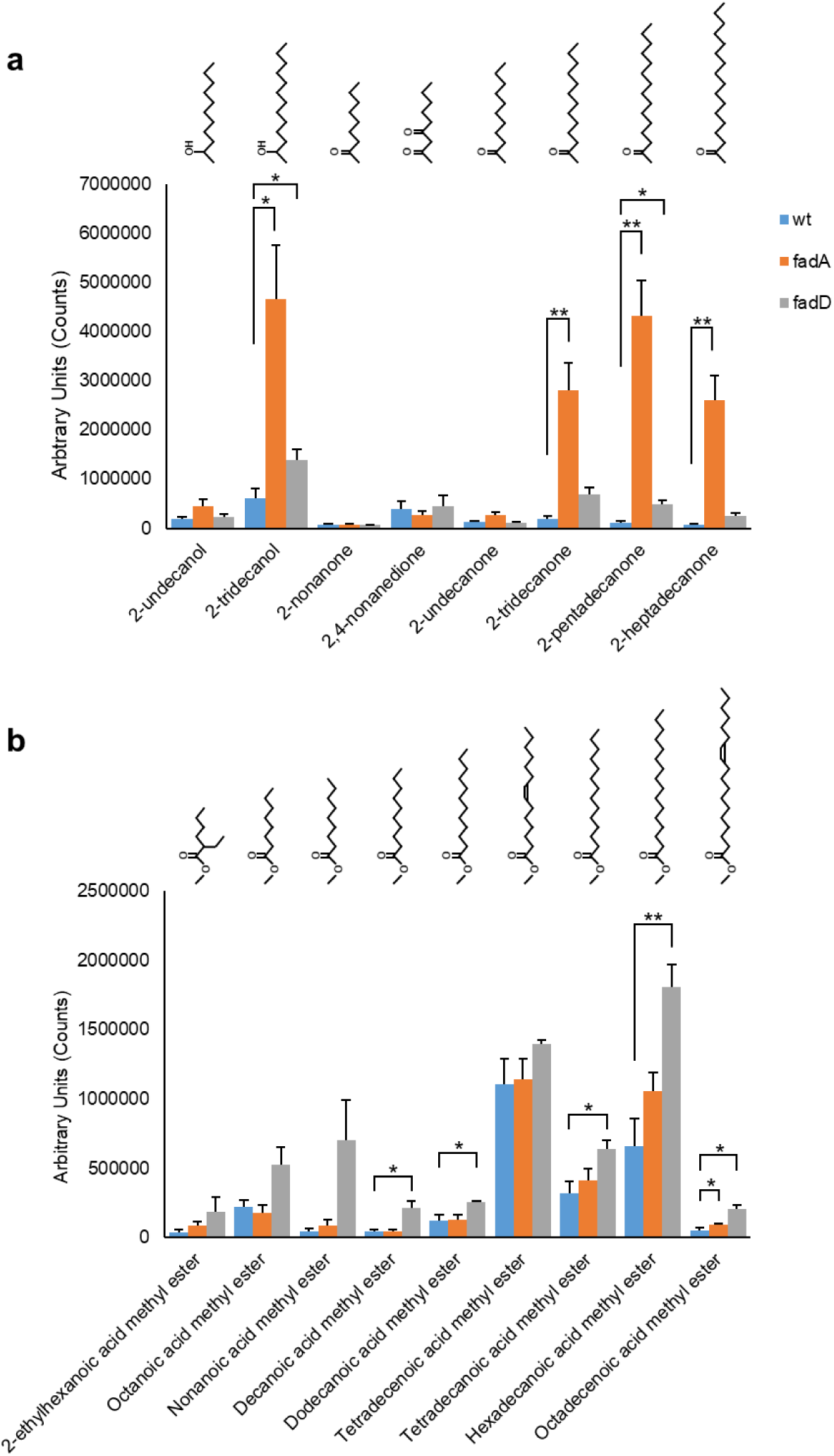
VOC profiles obtained by HS SPME-GC/MS of *S. meliloti* GR4 wild type (wt), and its *fadA* and the *fadD* mutant derivatives upon growth on solid 1% MM for 48 h. a) Volatilome profile of alcohols and methylketones (MKs); b) Volatilome profile of fatty acid methyl esters (FAMEs). Means of at least 3 biological replicates are indicated. Corresponding chemical structures are indicated above each graph. The positions of the C=C double bonds in tetradecenoic acid methyl ester and in octadecenoic acid methyl ester are tentative. Error bars indicate standard error at 95% confidence. Horizontal bars indicate significant difference: * p<0.05, ** p<0.01.

### Changes in bacterial communities in the soil exposed to *S. meliloti* volatilomes and synthetic volatile MKs

To determine the response of bacterial communities in the soil, DNA extracted from three biological replicates for each treatment (Sm volatilomes or synthetic MKs) at the initial time point (T=0), or after 1 or 7 days of exposition were subjected to amplicon sequencing of the V3-V4 region of the 16S rRNA gene. For alpha diversity in the volatilome experiment, after removing an outlier replicate from the 7-day control samples, the initial control (uninoculated MM plates at T=0) did not show significant differences in bacterial diversity (Shannon index Fig. 2a) or richness (observed ASVs Fig. 2a) with any of the other conditions or times. The samples with the highest diversity and richness were those exposed to the volatilome of the *fadA* mutant after 7 days. The only samples that showed significant differences compared to these were either the control samples after 1 day or those exposed for 1 day to the volatilome of the wild type, both of which displayed lower diversity and richness. More variability was observed in the alpha diversity indices of soil samples exposed to volatile MKs. In these experiments, the diversity and richness indices of the initial control were statistically different (p<0.05) from those of the control after 1 day but not after 7 days. Compared to their respective controls, 1 day of exposure to 2-TDC or the MIX (MK mix consisting of 2-TDC, 2-PDC and 2 HpDC) significantly increased diversity and richness, whereas after 7 days, exposure to the MK mix led to a significant decrease in both diversity and richness. In summary, the strongest effect observed in the soil experiment was in the diversity and richness of soil bacterial communities exposed to the MK mix which increased significantly after 1 day and decreased significantly after 7 days.

**Fig. 2.**
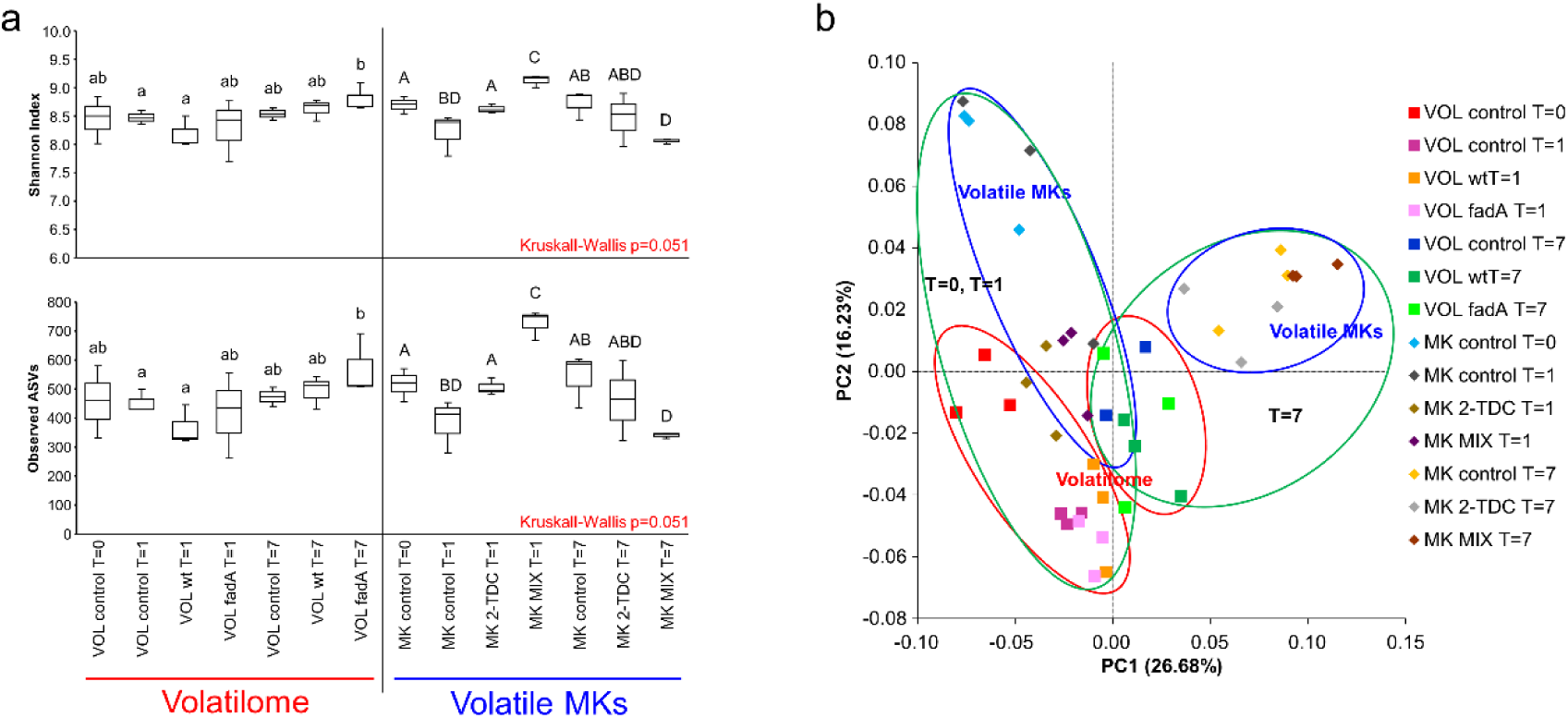
Effect of exposure to Sm volatilomes (VOL) and volatile MKs (MK) on the alpha and beta diversity of soil bacterial communities. a) Box and whisker plots of the Shannon Index for bacterial diversity and of the observed ASVs for bacterial richness. Different letters indicate significant differences (p<0.05) according to pairwise Kruskal-Wallis analysis. MIX indicates the mixture of 2-TDC, 2-PDC and 2-HpDC. b) PCoA analysis of samples from the soil assays based on weighted UniFrac distances. Color schemes and symbols are indicated on the right of the figure. Circles indicate clusters: green circles group samples of the time periods T=0 and T=1 together or of the time period T=7; blue circles subcluster volatile MKs samples and red circles subcluster Sm volatilomes samples.

To determine the effect of Sm volatilomes and volatile MKs on soil bacterial community structures, beta diversity was studied using principal coordinates analysis (PCoA) based on weighted Unifrac distances (Fig. 2b). The groupings show that the major effect was according to time (PERMANOVA pseudo-F 2.72, p=0.001) especially after 7 days. The bacterial community structures were also affected by whether they had been exposed to Sm volatilomes or volatile MKs (PERMANOVA pseudo-F 2.64, p=0.001) and the type of treatments: volatilome controls against exposure to either the wild type or *fadA* mutant volatilomes, volatile MKs controls or exposure to volatile 2-TDC or the MK mix (PERMANOVA pseudo-F 1.38, p=0.001). However, when replicate samples were compared pairwise to other replicate samples no significant differences were detected.

The ASVs detected in replicate samples at the different time points and treatments of soil samples exposed to Sm volatilomes and volatile MKs were classified to the phylum level and their mean relative abundance studied (Fig. 3a). The most abundant phyla in soil were Proteobacteria, Acidobacteriota, Gemmatimonadota, Bacteroidota and Actinobacteriota (Fig 3a) Under control conditions important shifts were already observable after only 1 day of incubation in both experiments. Furthermore, the changes in relative abundance of some phyla under the different conditions did not evolve linearly along time. Often, an abrupt increase or decrease was observed after 1 day, indicating that the populations had not yet stabilized. Therefore, we focused on changes between the initial T=0 samples and after 7 days of exposition. In both the Sm volatilome and the volatile MK experiments, the abundance of Bacteroidota and Proteobacteria increased and Actinobacteriota decreased under control conditions over time. Exposition to either the wild type or *fadA* mutant volatilomes further reduced the abundance of Actinobacteriota. On the other hand, exposition to either 2-TDC or the MK mix had different effects compared to each other, with the MK mix appearing to have a similar impact as exposition to the Sm volatilomes in decreasing Actinobacteriota abundance.

**Fig. 3.**
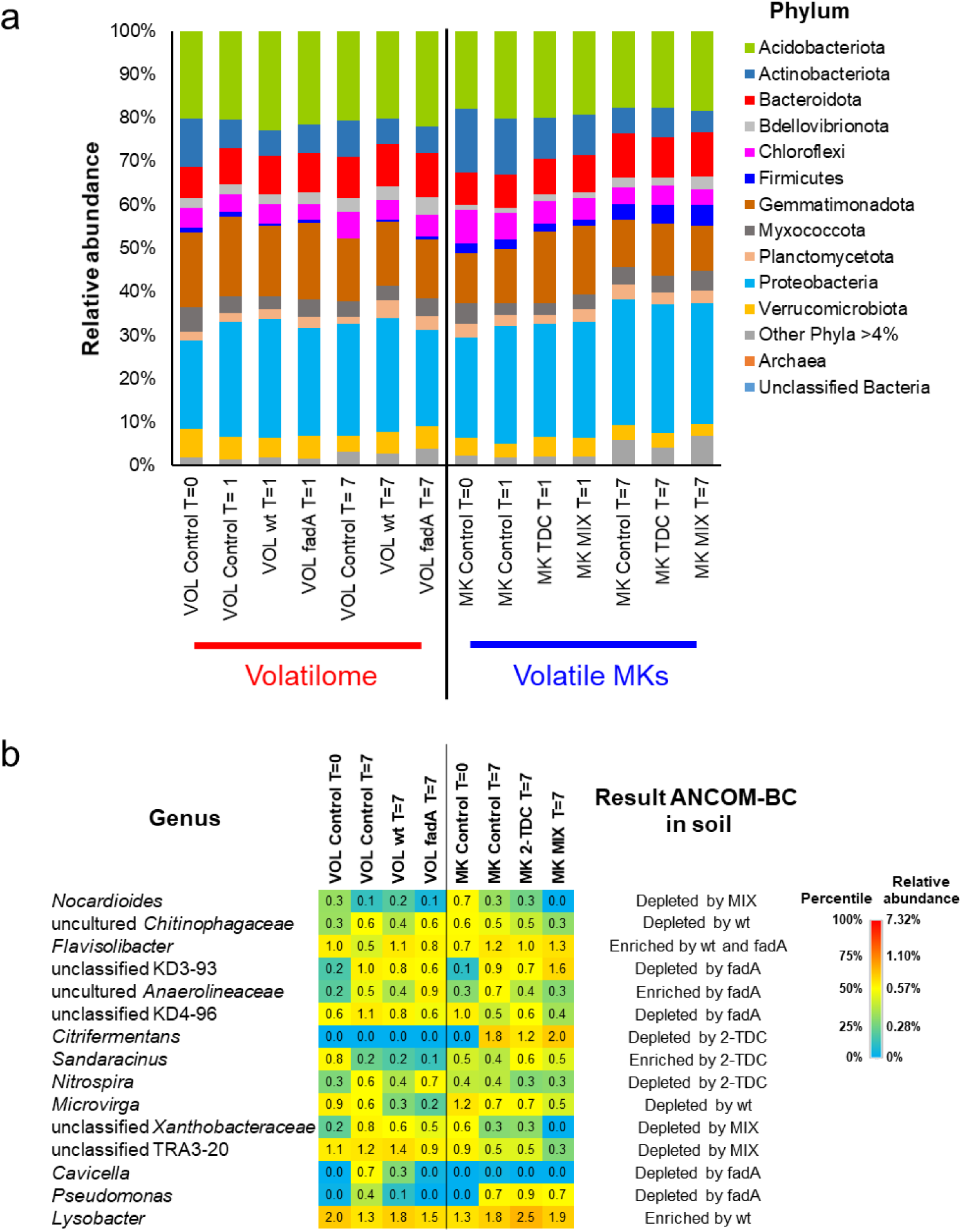
Identification of abundant taxa in soil assays. a) Mean relative abundances of the different bacterial phyla (> 4% abundance) of replicate samples for each condition and time in the soil assays. VOL for Sm volatilomes, MK for volatile methylketones, and MIX for the mixture of 2-TDC, 2-PDC and 2-HpDC. b) Heatmap of mean relative abundances of abundant bacterial genera in soil detected to be differentially abundant by ANCOM-BC (q-value < 0.05) compared to controls after 7 days within each treatment (exposure to Sm volatilomes (VOL) or to volatile methylketones (MK)). Heatmap data and color scale are based on percentiles extracted from the heatmap of the 90 most abundant genera in Fig. S3.

A closer look at the most abundant genera found in soil exposed to Sm volatilomes and volatile MKs soil (Fig. S3) revealed that at the initial time point (T=0) their relative abundances often differed between each experiment indicating that even using the same soil some changes occurred in bacterial populations during storage or the time past between experiments. As was also observed at the phylum level, for both experiments and under all treatments including controls, major shifts of abundant genera were seen after only 1 day of incubation. Since these changes were probably due more to perturbations in the bacterial populations caused by the setting up of the experiment (adjusting to humidity, oxygenation by physical mixing, etc) rather than by the different treatments used, the ANCOM-BC global test (Lin and Peddada 2020) was used to, principally, identify differentially abundant genera between the initial control and the different treatments after 7 days of incubation (Fig. 3b). Exposure of soil to the volatilome of the wild type enriched the bacteroidetal genus *Flavisolibacter* and the proteobacterial genus *Lysobacter* in soil (q-value < 0.05) and depleted the uncultured *Chitinophagaceae* and the proteobacterial genus *Microvirga* (q-value < 0.05). *Microvirga* abundance decreased even more upon exposure to the *fadA* mutant volatilome but not significantly (q-value > 0.05) (Fig. 3b). In the case of exposure to the volatilome of the *fadA* mutant, again *Flavisolibacter* was enriched as well as the Chloroflexi uncultured *Anaerolineaceae* (q-value < 0.05). On the other hand, exposure to the volatilome of this MK overproducing strain caused a decrease of the Chloroflexi genera KD4-96, the bacteroidetal genus *Sphingobacteriales* KD3-93 and the proteobacterial genera *Cavicella* and *Pseudomonas* (q-value < 0.05). For both proteobacterial genera, the small recuperation of *Cavicella* and *Pseudomonas* in the control after 7 days was hardly observed when exposed to the wild type volatilome. In the volatile MK experiment, when the soil communities were exposed to volatile 2-TDC, the abundance of *Nitrospira* decreased (q-value < 0.05) and the desulfobacterial genus *Citrifermentans* did not increase as much (q-value < 0.05) as in the control conditions after the same period of time. The myxococcal genus *Sandaracinus* only showed a significant increase in abundance when exposed to volatile 2-TDC (q-value < 0.05). Exposure to the MK mixture decreased (q-value < 0.05) the abundance of ASVs classified as the actinobacterial genus *Nocardioides*, an unclassified genus belonging to the proteobacterial family *Xanthobacteraceae* and an unclassified burkholderiales genus belonging to TRA3-20. The genus *Ensifer* is synonymous to *Sinorhizobium* (Fagorzi et al. 2020) so in bacterial population analyses both will be referred to as *Ensifer*/*Sinorhizobium*. *Ensifer*/*Sinorhizobium* were amongst the 90 most abundant genera in the soil assays (Fig. S3). This taxon did not give significant differences with ANCOM-BC but did respond differently in relative abundance in the Sm volatilome experiment compared to the volatile MK experiment where in the former it almost disappeared after 7 days without showing significant differences between times or treatments (Fig. S4). On the other hand, in the MK experiment a significant (p<0.05) increase could be observed in soil exposed to 2-TDC or the MK mixture after 7 days compared to initial (T=0) control conditions. For the MK mixture this increase was also significant when compared to control conditions after 7 days (Fig. S4).

### Changes in plant associated bacterial communities exposed to *S. meliloti* volatilomes and synthetic volatile MKs

The response of bacterial communities in the rhizosphere and the root endosphere of *Medicago truncatula* to exposition to the volatilomes of the wild type or the *fadA* mutant or to volatile 2-TDC or the MK mixture were determined by 16S rRNA amplicon sequencing. The DNA used was extracted from three biological replicates for each treatment at the initial time point and after 14 days of exposition. In the rhizosphere samples, the alpha diversity indices of all the samples together were significantly higher than of all the root endosphere samples (p< 10^-9^, Fig. S5). However, within the rhizosphere samples no significant differences were observed in Shannon diversity or richness (Fig. 4a). On the other hand, in the root endosphere samples, after the removal of an outlier replicate from the endosphere samples exposed to the wild type volatilome after 14 days, alpha diversity indices indicated significant differences (p<0.05) in diversity and richness (Fig. 4b). As expected, the values for diversity and richness of the initial (T=0) control endosphere samples of either the Sm volatilome or the volatile MKs experiments were similar (p> 0.05) to each other. Likewise, the diversity and richness of the controls after 14 days from each experiment were similar to each other but diversity was significantly higher (p< 0.05) than the initial (T=0) controls (Fig. 4b). The diversity or richness of those communities exposed to the volatilomes of the wild type or the *fadA* mutant after 14 days however did not change significantly from the initial T=0 time point while the diversity or richness of those exposed to 2-TDC or the MK mixture were as high as the controls after 14 days (Fig. 4b). Therefore, the most dramatic effect observed was that exposure to Sm volatilomes maintained the initial low diversity in the root endosphere bacterial populations while controls or exposure to synthetic volatile MKs led to an increase of diversity over time.

**Fig. 4.**
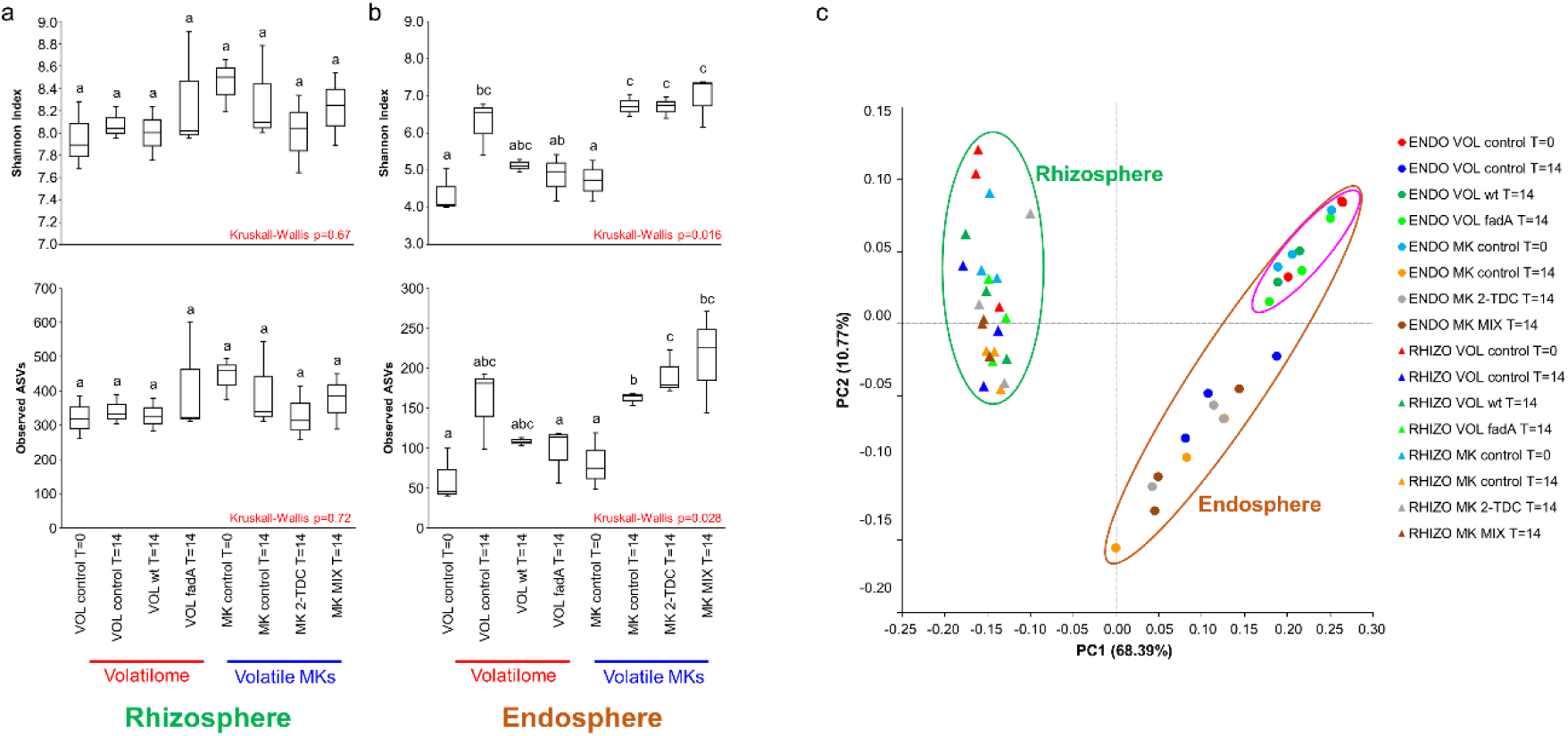
Effect of exposure to Sm volatilomes (VOL) and volatile MKs (MK) on the alpha and beta diversity of the bacterial communities associated with *M. truncatula* plants. a) Box and whisker plots of the Shannon Index for bacterial diversity and of the observed ASVs for bacterial richness in the rhizosphere. b) Box and whisker plots of the Shannon Index for bacterial diversity and of the observed ASVs for bacterial richness in the root endosphere. Different letters indicate significant differences (p<0.05) according to pairwise Kruskal-Wallis analysis. MIX indicates the mixture of 2-TDC, 2-PDC and 2-HpDC. c) PCoA analysis of the samples from the plant assays based on weighted UniFrac distances. Color schemes and symbols are indicated on the right of the figure. Circles indicate clusters: green circle groups rhizosphere samples; brown circle groups root endosphere samples; pink circle groups endosphere Sm volatilome T=0, volatile MKs T=0 and Sm volatilomes T=14 samples.

To determine the effect of Sm volatilomes and synthetic volatile MKs on the rhizosphere or root endosphere bacterial community structures, beta diversity was studied by PCoA (Fig. 4c). The groupings show that the major effect was according to sample type, i.e. whether the bacterial communities were found in the rhizosphere or the root endosphere (PERMANOVA pseudo-F 74.54, p= 0.001). The other factors did not have significant effects on their own. However, the combination of sample type and time had a significant interaction (ADONIS p=0.021) as well as the interaction between sample type and experiment (Sm volatilome vs volatile MKs, ADONIS p=0.026). Those communities in the root endosphere which shared low diversity or richness (Fig. 4b; initial T=0 controls as well as communities exposed to the volatilomes of the wild type or *fadA* mutant) clustered together (Fig. 4c: red circle).

According to the taxonomic identities of the ASVs detected in the rhizosphere soil of plants exposed to Sm volatilomes or synthetic volatile MKs the most abundant phyla were Proteobacteria followed by Bacteroidota, Firmicutes, Acidobacteriota, Actinobacteriota, Chloroflexi, Myxococcota and Verrucomicrobiota (Fig. 5a). However, the changes observed within these phyla between treatments and times in either the Sm volatilome or volatile MK experiments were small and mostly not significant. On the other hand, fewer abundant phyla were observed in the root endosphere samples, with Proteobacteria predominating followed by Firmicutes, Bacteroidota and Actinobacteriota (Fig. 5a). Compared to the initial populations, control root endosphere samples after 14 days showed a marked and significant decrease in the relative abundance of Proteobacteria and an increase in Firmicutes (one-tailed T test, p < 0.05) in both the Sm volatilome and the volatile MKs experiments. Interestingly, the relative abundances of these phyla in the root endosphere of plants exposed to the volatilome of the wild type or *fadA* mutant were not significantly different from T=0 controls but were significantly different (p < 0.05) from the 14-day controls of either experiment or in the root endosphere of volatile MK exposed plants. On the other hand, in the volatile MK experiment, the phylum profile in the root endosphere of plants exposed to 2-TDC or the MK mixture were more similar to the T=14 controls than the initial T=0 controls. Altogether, in the root endosphere, the relative abundance of the most abundant phyla in the initial and Sm volatilome-exposed plants after 14 days were more similar to each other than to those in the controls after 14 days and volatile MK-exposed plants.

**Fig. 5.**
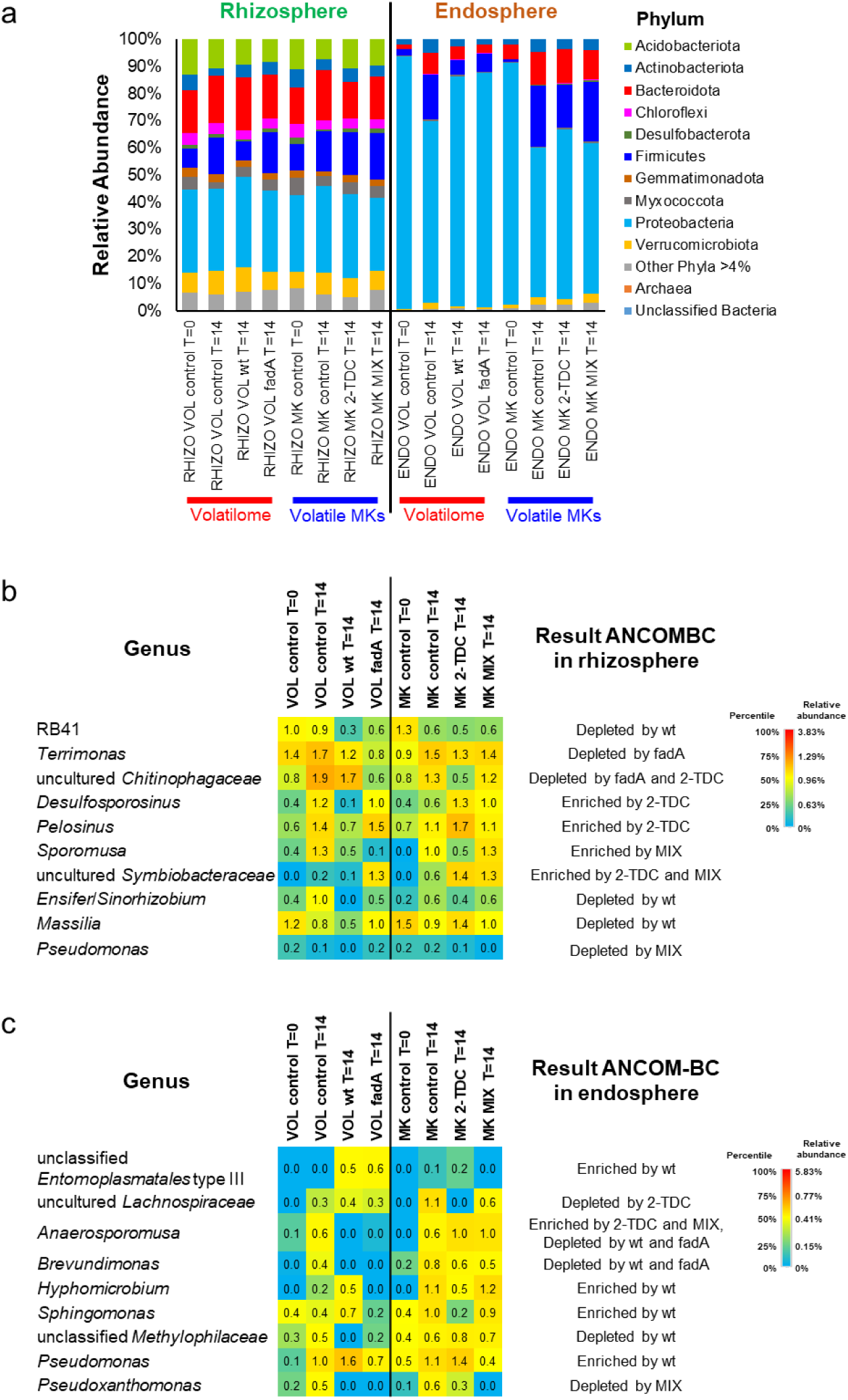
Identification of abundant taxa in plant assays. a) Mean relative abundances of the different bacterial phyla (> 4% abundance) of replicate samples for each condition and time in the plant assays. RHIZO for rhizosphere, ENDO indicates root endosphere, VOL for Sm volatilomes, MK for volatile MKs, and MIX for the mixture of 2-TDC, 2-PDC and 2-HpDC. b) Heatmap of mean relative abundances of abundant bacterial genera in the rhizosphere detected to be differentially abundant by ANCOM-BC (q-value < 0.05) compared to controls after 14 days within each treatment (exposure to Sm volatilomes (VOL) or to volatile MKs (MK)). Heatmap data (% rel. abundance) and color scale based on percentiles extracted from the 49 most abundant rhizosphere bacterial genera in Fig. S6. c) Heatmap of the mean relative abundances of abundant bacterial genera in the root endosphere excluding *Ensifer*/*Sinorhizobium* which were detected to be differentially abundant by ANCOM-BC (q-value < 0.05) compared to controls after 14 days within each treatment. Heatmap data (% rel. abundance) and color scale based on percentiles extracted from the 53 most abundant root endosphere bacterial genera excluding *Ensifer*/*Sinorhizobium* in Fig. S7.

In order to further determine how the most abundant genera were affected by exposure to volatilomes or volatile MKs in the rhizosphere or root endosphere of plants, the relative abundance of each abundant genus was analyzed with heatmaps (Figs. S6, S7, respectively). Furthermore, differentially abundant genera analyzed by the ANCOM-BC global test (Lin and Peddada 2020) showed that in the rhizosphere (Fig. 5b) in response to exposure to the volatilome of the wild type, the acidobacterial genus RB41, and the proteobacterial genera *Massilia* and *Ensifer*/*Sinorhizobium* were significantly depleted (q < 0.05) compared to controls after 14 days. In response to exposure to the volatilome of the *fadA* mutant, the bacteroidetal genera *Terrimonas* and uncultured *Chitinophagaceae* were depleted (q < 0.05). No abundant genera were detected which were significantly enriched by the presence of Sm volatilomes. Uncultured *Chitinophagaceae* was also depleted (q < 0.05) upon exposure to 2-TDC while the Firmicutes genera *Desulfosporosinus*, *Pelosinus*, and uncultured *Symbiobacteraceae* were enriched (q < 0.05). Uncultured *Symbiobacteraceae* was also enriched in the presence of the MK mixture (q < 0.05) and showed increased abundance with the volatilome of the *fadA* mutant (not significant). The Firmicutes genus *Sporomusa* showed differential abundance in response to the MK mix in which it was enriched (q < 0.05). *Pseudomonas* was not amongst the most abundant genera in the rhizosphere but it was included in the heatmap (Fig. S6) as it was also depleted (q < 0.05) in the presence of the MK mixture compared to controls after 14 days (Fig. 5b). In general, when a significant differential response could be detected in abundant rhizosphere genera, exposure to Sm volatilomes mostly had a deleterious effect while volatile MKs had an enriching effect.

In the root endosphere most of the abundant genera excluding *Ensifer*/*Sinorhizobium* had very low abundance initially and increased after 14 days in the controls (Fig. S7). Using the ANCOM-BC global test to find differentially abundant genera by comparing each condition to controls after 14 days within each treatment (Sm volatilome or volatile MKs) (Fig. 5c) revealed that Firmicutes genus *Anaerosporomusa*, the proteobacterial genus *Brevundimonas* and an unclassified genus belonging to the proteobacterial family *Methylophilaceae,* were depleted under exposure to the volatilome of the wild type (q- value < 0.05). Of these, *Anaerosporomusa* and *Brevundimonas* were also depleted under exposure to the volatilome of the *fadA* mutant. Curiously, *Anaerosporomusa* was enriched with 2-TDC and the MK mixture (q < 0.05), while the other two taxa maintained similar abundance levels with the MKs. In contrast to the above mentioned taxa, an unclassified genus belonging to the Firmicutes family *Entomoplasmatales* type III, and the proteobacterial genera *Hyphomicrobium*, *Sphingomonas*, and *Pseudomonas* were enriched by exposure to the wild type volatilome (q<0.05). With regard to exposure to synthetic volatile MKs, uncultured *Lachnospiraceae* and the proteobacterial genera *Pseudoxanthomonas* were depleted (q < 0.05) in the presence of 2-TDC and with the MK mixture, respectively (Fig. 5c). For both taxa, their relative abundance was either unaffected (uncultured *Lachnospiraceae*) or decreased (*Pseudoxanthomonas*) by exposure to the volatilomes of the wild type or the *fadA* mutant (not significant).

As *Ensifer*/*Sinorhizobium* are the natural symbionts of *M. truncatula*, a closer look at these populations at the level of genera was taken in the rhizosphere and the root endosphere (Fig 6). In the rhizosphere, the relative abundances of *Ensifer*/*Sinorhizobium* were only approximately 0.5 % (Fig 6a). In the Sm volatilome experiment, a small but not significant increase was observed in *Ensifer*/*Sinorhizobium* abundance in the in the rhizosphere of control plants after 14 days compared to initial relative abundances. In the rhizosphere of plants exposed to the wild type volatilome, *Ensifer*/*Sinorhizobium* became undetectable after 14 days (Fig 6a) which constituted a significantly (p<0.05) lower abundance compared to the control after 14 days but not with the initial (T=0) control. In contrast, no effect due to synthetic MKs or time was observed on *Ensifer*/*Sinorhizobium* abundance in the rhizosphere of the volatile MK experiment (Fig 6a). On the other hand, the root endosphere was dominated by *Ensifer*/*Sinorhizobium* (Fig. 6b). The high proportion of *Ensifer*/*Sinorhizobium* was provided by only 26 ASVs, especially by 10, which accounted for 92% of all reads identified to be *Ensifer*/*Sinorhizobium*. The high abundance of *Ensifer*/*Sinorhizobium* was expected as the *M. truncatula* plants exhibited nodules but the number of nodules per plant was not scored in these experiments. Neither were nodule microbial populations determined separately from root endophytic populations but studied together as the whole root endosphere. The highly abundant *Ensifer*/*Sinorhizobium* in the endosphere shows the same pattern as Proteobacteria detected at the phylum level in response to time and treatment in which, once more, the lowering of *Ensifer*/*Sinorhizobium* abundance in the control samples after 14 days in both the Sm volatilome and the volatile MK experiments was countered by the exposition to the volatilomes of the wild type and the *fadA* mutant but not by exposure to 2-TDC or the MK mixture

**Fig. 6.**
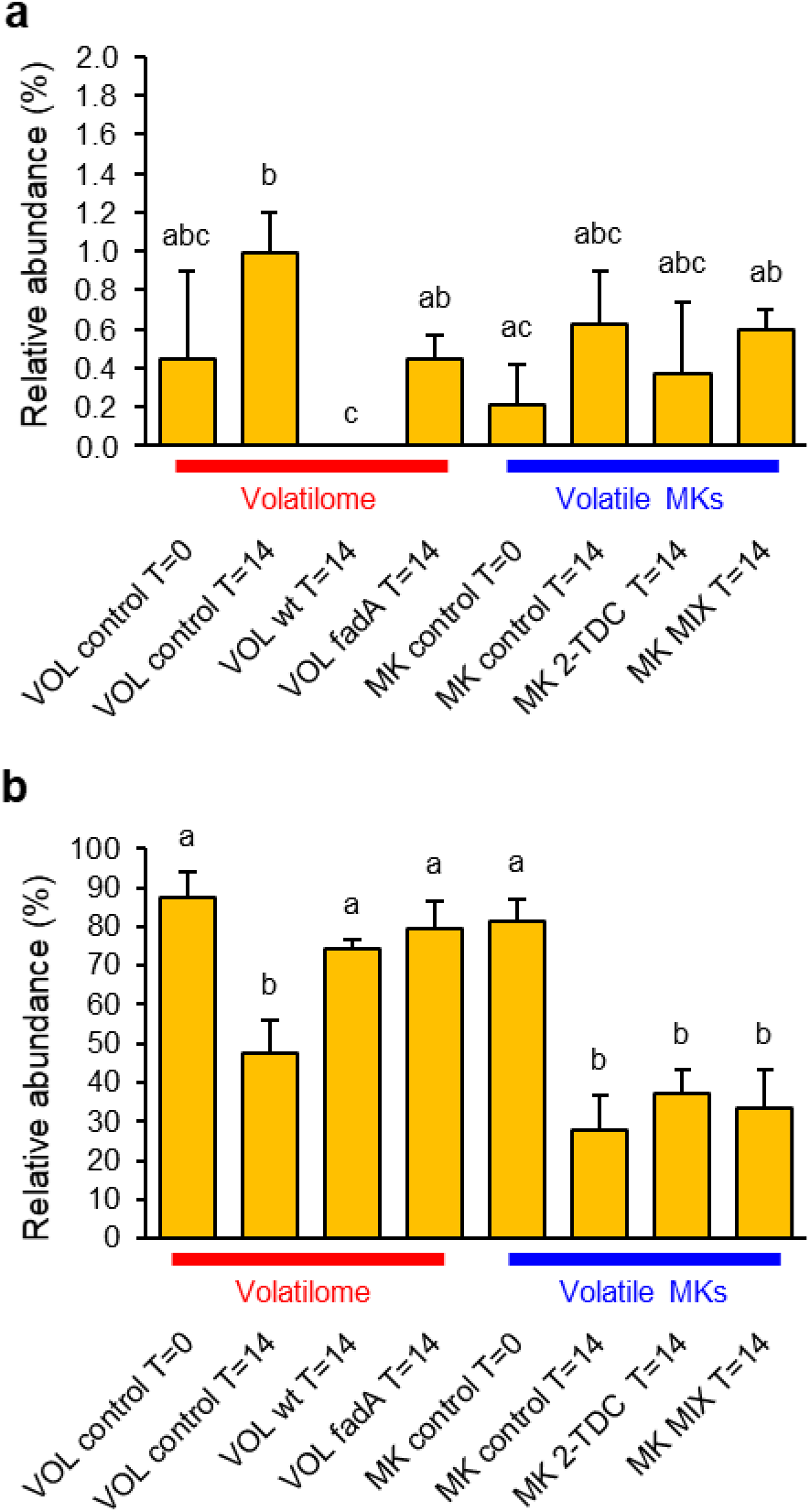
Relative abundance of *Ensifer*/*Sinorhizobium* in the a) rhizosphere and b) root endosphere of *M. truncatula* plants exposed to Sm volatilomes (VOL) or volatile MKs (MK). MIX indicates the mixture of 2-TDC, 2-PDC and 2-HpDC. Error bars indicate standard error. Different letters indicate significant differences (Welch T-test; p<0.05).

## Discussion

### *S. meliloti* volatilomes comprise fatty acid-derivative compounds

Analyses of the volatile organic compounds released by the wild type *S. meliloti* GR4 and its *fadD*, or *fadA* mutant derivatives led to the identification of seventeen fatty acid-derivative compounds comprising methylketones (MKs), alcohols, and fatty acid methyl esters (FAMEs). The production of the MKs 2-tridecanone (2-TDC), 2-pentadecanone (2-PDC) and 2-heptadecanone (2-HpDC) was confirmed to be much higher in the *fadA* mutant than in the wild type strain, or in the *fadD* mutant, which had previously been shown to emit higher levels of the three MKs than the wild type (López-Lara et al. 2018). These results help to understand how MKs are produced in Sm by suggesting that they can be derived not only from intermediates of the fatty acid biosynthesis pathway i.e. 3- oxo-acyl-ACP as proposed by López-Lara et al. (2018) but also from fatty acid degradation (β-oxidation) intermediates, namely the FadA substrate 3-oxo-acyl-CoA (Fig. S1). Recently, the YbgC-like thioesterase SMc03960 of Sm was described as an acyl-ACP/acyl-CoA thioesterase with broad substrate specificity that contributes to 2- TDC formation (Bernabéu-Roda et al. 2025). Nonetheless, the participation of SMc03960 in the increased production of MKs by the *fadA* mutant remains to be investigated.

Of the two alcohols detected in the VOC profiles of the three Sm strains analyzed, the emission of 2-tridecanol reflects the same patterns as observed for 2-TDC production, i.e. very high in the *fadA* mutant, high in the *fadD* mutant, and low in the wild type strain. This suggests that this methylalcohol may derive from the reduction of 2-TDC by, for instance, an alcohol dehydrogenase (Fig. S1). Whereas no biological activity has been reported yet for 2-tridecanol, 2-undecanol has been shown to inhibit to some degree phytopathogenic fungi (Freitas et al. 2022; Wu et al. 2019; Yuan et al. 2012).

The production of up to 9 FAMEs and in high quantities by Sm is noteworthy. Although FAMEs have been identified in bacteria, especially as technology has improved for VOC detection, their presence is usually limited to a few FAMEs and in small amounts compared to other VCs (Weete et al. 1970, Flavier et al. 1997; Fitzgerald et al. 2020; Wakimoto et al. 2020; Zaid et al. 2023; Ren et al. 2024). Two long-chain FAMEs, methyl 3-hydroxypalmitate (3-OH-PAME) and methyl 3-hydroxymyristate (3-OH-MAME), have been identified as diffusible volatile factors that function as autoinducers of the *phc* quorum sensing system in *Ralstonia solanacearum* (Flavier et al. 1997; Kai 2023), and in *Cupriavidus taiwanensis* (Wakimoto et al. 2020). In these bacteria, these FAMEs are produced by the action of PhcB, a class I *S*-adenosylmethionine (SAM)-dependent *O*-methyltransferase, which catalyses the SAM-dependent methylation of the free fatty acids 3-OH PA and 3-OH MA, respectively (Ujita et al. 2019). Since increased amounts of FAMEs are produced by the *fadD* mutant, which is known to accumulate free fatty acids (Pech-Canul et al. 2011), the action of as yet unidentified SAM-dependent methyltransferases are probably responsible for their production in Sm (Fig. S1). Besides a role in quorum sensing regulation, different FAMES have been recently shown to inhibit the growth of pathogenic fungi or influence bacterial motility and biofilm formation (Zaid et al. 2023; Ren et al. 2024). This highlights that the volatilome emitted by *S. meliloti* GR4 contains compounds, in addition to the MKs, which may have biological activity with an impact on soil and plant associated microbiota.

### Effect of *S. meliloti* volatilomes and synthetic volatile MKs on soil bacterial communities

The aim of the current study was to determine the ecological role of MKs on natural bacterial populations by determining the effect of Sm volatilomes with different MK content or synthetic MKs on soil and plant microbiomes. In the soil assays, both alpha- and beta-diversity were generally little affected by Sm volatilomes or synthetic MKs and the major changes in the most abundant phyla were variable. However, exposure to Sm wild type and *fadA* mutant volatilomes as well as to the MK mixture coincided in decreasing the abundance of Actinobacteriota, a phylum that includes beneficial plant interacting bacteria and bacteria that produce important secondary metabolites such as antibiotics. In the study by Yuan et al. (2017), exposition of soil bacterial communities to the volatilome of *B. amyloliquefaciens* NJN-6 led to a significantly lower diversity, altered the bacterial community structure and, as in our study, downward shifts of the abundance of Actinobacteriota. Interestingly, the volatilome profile of *B. amyloliquefaciens* NJN-6 is characterized by high levels of MKs (Yuan et al. 2012), which is compatible with a putative role of MKs on the observed effect on Actinobacteriota abundance.

A number of abundant genera were identified in the soil to be differentially affected by exposition to Sm volatilomes or synthetic MKs. Among the genera stimulated by exposure to Sm volatilomes and that have been related to legumes include *Flavisolibacter*, a genus that has been associated with alfalfa yield (Omirou et al. 2024), which was enriched by both the wild type and *fadA* mutant volatilomes but unaffected by synthetic MKs. *Lysobacter*, a known antifungal biocontrol bacterium, with possible roles for the treatment of plant diseases and for plant growth promotion (Lin et al. 2021), and which has previously been observed in bulk soil as well as in the rhizosphere, rhizoplane, phyllosphere, roots and nodules of *M. truncatula* (Tkacz et al. 2020) was also stimulated by exposure to the wild type volatilome in our study. On the other hand, *Ensifer/Sinorhizobium* was found to increase its abundance upon exposure to synthetic MKs but did not become significantly enriched. Among the genera which became depleted upon exposure to Sm volatilomes included *Microvirga*, a genus which has been described to induce nodule formation (Martínez-Hidalgo and Hirsch 2017) and to be associated with alfalfa yield (Omirou et al. 2024). *Pseudomonas*, a highly diverse, ubiquitous and metabolic versatile genus which includes plant beneficial and pathogenic species (Peix et al. 2018), was also observed to be depleted in soil exposed to Sm volatilomes but did not respond to synthetic MKs. This observation is contrary to that described by Baroja-Fernandez et al. (2021), in which *Pseudomonas* was enriched in soil amended with fungal cell free extracts or distillates. Further investigation will be needed to determine how and why the affected genera respond as they do and to which components of the Sm volatilomes. Moreover, which species within these genera are sensitive to these components and which functions they play in the soil microbiome remain open questions.

### Effect of *S. meliloti* volatilomes and synthetic volatile MKs on plant associated bacterial communities

This work represents the first study of the effect of bacterial volatilomes or bacterial organic volatiles as airborne compounds on plant associated microbiomes. In the bacterial communities of the rhizosphere of *M. truncatula*, exposure to Sm volatilomes or volatile MKs did not have clear effects on alpha or beta diversity and on the most abundant phyla, the effect was variable. When differential responses were determined within the most abundant rhizosphere genera, of those which showed a significant response exposure to Sm volatilomes mostly had a depleting effect while volatile MKs had an enriching effect. For example, RB41, a bacteroidetal genus found in the rhizosphere of *M. truncatula* which responds to plant-derived organic matter (Liu et al. 2023), *Massilia* which has been found in the rhizosphere of soybean (Han et al. 2024) and *M. truncatula* (Tkacz et al. 2020) and to be enriched in soil planted with pepper and amended with cell-free culture filtrates and distillates (Baroja-Fernandez et al. 2021), and *Ensifer/Sinorhizobium* were all depleted in the rhizosphere by exposition to wild type volatilomes. Uncultured *Chitinophagaceae*, a representative of a family found in the rhizosphere of soybean (Han et al. 2024), is an example of a taxon that became depleted in the rhizosphere of plants exposed to the *fadA* mutant volatilome but also to 2-TDC and therefore could represent a taxon that shows sensitivity to MKs. Within the abundant genera which became enriched upon exposition to volatile MKs all belong to Firmicutes which have not previously been associated with plants. However, one taxon, uncultured *Symbiobacteraceae*, a member of a family correlated with soil N_2_O emissions (Hao et al. 2022) not only became enriched with volatile MKs but also increased its abundance, although not significantly, upon exposure to the volatilome of the *fadA* mutant. This suggests that this taxon may be responsive to higher MK concentrations in the rhizosphere. On the other hand, *Pseudomonas* was curiously shown to be depleted in the rhizosphere by the MK mixture. Besides a direct role of bacterial VOCs on microbial traits that can impact interactions with plants (Schulz-Bohm et al. 2017; López-Lara et al. 2018; Sharifi et al. 2022; Weisskopf et al. 2021), VOCs may also affect plant growth and exudate composition (Hernandez-Calderon et al 2018; Almeida et al 2022). Therefore, the shifts observed for specific genera in the rhizosphere of *M. truncatula* plants exposed to Sm volatilomes or volatile MKs in our experiments could be due not only to a direct effect on specific microbiome members but also as an indirect effect on the plant to produce different organic compounds in their exudates with nutritional, attractant or growth inhibiting/ defensive characteristics leading to the selection of specific bacterial genera.

The clearest effects of Sm volatilomes were observed in the root endosphere where exposition to Sm volatilomes for 14 days resulted in the maintenance of the low initial (T=0) diversity values in the root endosphere of *M. truncatula* plants compared to control plants or those exposed to synthetic MKs, in which the diversity of root endosphere bacterial communities increased with time. The diversity of plant associated microbiota has been found to be lower in healthy plants than in plants faced with dysbiosis (Arnault, et al. 2023). Lower bacterial diversity was also observed in the endosphere of *M. truncatula* growing in rich nutrient soils than in poorer soils (Lagunas et al 2023). As bacterial populations approach the plant interior, they become more restrictive and subject to plant control (Bulgarelli et al 2013, Muller et al., 2016, Uribe-Acosta et al 2025). This suggests that plants actively select those bacteria which enter the endosphere, and therefore the bacterial diversity inside plant tissues could depend, at least in part, on how the plant perceives its health and nutritional status. In this manner it could be possible that the volatilomes or VCs emitted by a legume beneficial symbiont convey signals to the plant of wellbeing inducing the plant to recruit less additional bacterial species into its endosphere and thereby maintaining low diversity.

In the root endosphere the most abundant phylum corresponded to Proteobacteria which consisted mostly of *Ensifer*/*Sinorhizobium* which dominated originally and remained dominant in the presence of Sm volatilomes but declined with time in control plants and MK-exposed plant roots with a concomitant increase of Firmicutes and Bacteroidota relative abundance. Although a time effect was not studied, the root endosphere of *M. truncatula* plants exposed to DMHDA in the growth medium caused concentration dependent shifts of Firmicutes, alpha- and beta-proteobacteria and Actinobacteriota (Real-Sosa et al. 2022). When looking at some of the most abundant genera, besides *Ensifer/Sinorhizobium*, which were differentially affected in the endosphere, of those which increased their abundance in the root endosphere upon exposure to Sm volatilomes and over time, included an unclassified member of *Entomoplasmatales* Type III typically associated with arthropods (Sapountzis et al. 2015). This may suggest the presence of root-borne insects in the samples. On the other hand, the impact of Sm volatilomes had an opposite tendency than *Ensifer*/*Sinorhizobium* for *Brevundimonas*, which have been detected in the endosphere of various plants (Cheng et al. 2019; Ghiasvand et al. 2019), an unclassified genus belonging to *Methylophilaceae*, a family which have been found in the root and rhizosphere of lettuce (Persyn et al. 2024), and *Anaerosporomusa,* an anaerobic spore forming strain which also has been found as an endophyte in unhealthy potatoes (Rajapaksha et al. 2024), indicating that members of these genera are, somehow, restricted entrance into the root in the presence of Sm volatilomes. Curiously, *Anaerosporomusa*, became enriched with volatile MKs which could indicate this genus to be sensitive to other VCs present in the Sm volatilomes.

*Pseudomonas* also became enriched in the root endosphere in the presence of the wild type volatilome as well as with 2-TDC but its relative abundance declined in the presence of the *fadA* mutant volatilome or MK mixture. *Pseudomonas* has been described to be a common non-rhizobial endophyte (NREs) in nodules that can impact, in different ways the legume-rhizobium symbiosis (Yu et al. 2025; Kosmopoulos et al. 2024). Burns et al. (2021) identified the opportunistic pathogen *P. aeruginosa* (Walker et al. 2004), as well as plant growth promoting (PGP) species such as *P. fluorescens,* and *P. putida* in root nodules of *M. truncatula*. However, the identity at the species level of the *Pseudomonas* detected in our study and their specific location (inside or outside the nodules) remains unknown. 2-TDC has been shown to interfere with the capacity of pathogenic *Pseudomonas* species to form biofilms and colonize plant tissues without affecting growth (López-Lara et al. 2018) but its effect on their relative abundance within complex populations is unknown. Moreover, *Pseudomonas* responded differently in the endosphere than in the rhizosphere or in soil albeit in the latter case the incubation time and experimental set-up was different. This suggests that MKs or Sm volatilome components have niche-specific effects and probably species-specific effects within bacterial populations. Therefore, determining the species of *Pseudomonas* or of the other identified differentially affected genera, how and why they respond differently in the soil, rhizosphere and root endosphere to Sm volatilomes or MKs, and which roles they play for plant health merit further investigation.

The strongest effect observed in our experiments was that exposure to Sm volatilomes, but not to synthetic MKs, maintained root endophytic *Ensifer*/*Sinorhizobium* populations over time. These results suggest that component(s) other than MKs within the Sm volatilomes were responsible for this effect, unless this sensitivity was specific to a strict range of concentrations or exposure times. On the other hand, besides being much less abundant in the soil and rhizosphere, it appears that *Ensifer/Sinorhizobium* populations respond differently to Sm volatilomes and pure MKs in each niche. In the soil, *Ensifer/Sinorhizobium* populations were unaffected by exposure to Sm volatilomes but exposure to pure MKs increased their abundance especially upon exposition to the MK mixture. In contrast, Sm volatilome exposure in the rhizosphere especially with that emitted by the wild type strain resulted in lower abundance of *Ensifer/Sinorhizobium* while exposure to synthetic MKs had no effect. Therefore, it appears that volatile MKs affect *Ensifer/Sinorhizobium* populations only in the soil, while Sm volatilomes only affect *Ensifer/Sinorhizobium* populations under the influence of the plant by decreasing their abundance in the rhizosphere and increasing them in the endosphere. As bacterial VOCs have been shown to affect plant root exudate composition (Hernández-Calderón et al. 2018) on the one hand, and the plant controls bacterial as well as rhizobial infection, nodule formation and senescence, on the other, suggests that Sm volatilomes may also exert their influence on *Ensifer/Sinorhizobium* populations in the rhizosphere and the root endosphere through the plant. However, any effect of MKs on these populations could be overshadowed simply by the stronger influence of the plant itself. More studies will be needed to determine how MKs or Sm volatilome components affect the plant, but also how they directly influence *Ensifer/Sinorhizobium* populations and those of the other sensitive genera identified in the soil, rhizosphere and root endosphere.

## Conclusions

In this study we demonstrate, for the first time, the effect of *S. meliloti* volatilomes and airborne synthetic methylketones (MKs) on soil and plant microbiomes. Our major finding is that Sm volatilomes have a strong effect on maintaining high *Ensifer/Sinorhizobium* abundance in the endosphere of *M. truncatula* roots and hinder the natural tendency of additional different bacterial species to enter the root over time. In order to further advance our understanding of the effect of MKs on natural bacterial populations, mutants of fatty acid degradation pathways were studied. Mutation of *fadA* was found to overproduce MKs while inactivation of *fadD* led to an increase in the emission of fatty acid methyl esters (FAMEs). Thereby, the creation of the MK overproducing strain and the characterization of the volatilome profile of these mutants and the wild type strain has given new insights on the synthesis of rhizobial volatile compounds. Besides the important effect on *Ensifer/Sinorhizobium* populations in the root endosphere, members of the soil and plant associated microbiomes which are specifically affected by Sm emitted volatilomes and synthetic MKs were identified. Future investigation will be needed to determine to what extent these genera are affected by MKs and Sm volatilome components and what their functions are in the soil and plant associated microbiomes. This work constitutes, to the best of our knowledge, the first observation of how the volatilome emitted by Sm or any other bacteria affects plant microbiomes. Altogether, this study extends our knowledge of how bacterial volatilomes play a relevant role in communication in plant-bacteria interactions which could lead to new applications for sustainable agriculture.

## Acknowledgements

This work was supported by grant PID2021-123540NB-I00 funded by the Spanish Ministerio de Ciencia, Innovación y Universidades (MICIU) and the Spanish Agencia Estatal de Investigación (AEI) MICIU/AEI/10.13039/501100011033 with co-financing from the European Regional Development Fund “ERDF A way of making Europe” and by grant P20 00225 funded by the Consejería de Economía, Conocimiento, Empresas y Universidad (Junta de Andalucía, PAIDI)). This work was also supported by grant This work was also supported by UNAM-PAPIIT IN221525 funded by the Universidad Nacional Autónoma de México (UNAM). SPME analyses were carried out at the Instrumental Technical Services of the Estación Experimental del Zaidín (CSIC), Granada, Spain. We thank Ángeles Moreno-Ocampo for skillful technical assistance

## Statements and Declarations

### Funding

This work was supported by grant P20 00225 funded by the Consejería de Economía, Conocimiento, Empresas y Universidad (Junta de Andalucía, PAIDI)) and by grant PID2021-123540NB-I00 funded by the Spanish Ministerio de Ciencia, Innovación y Universidades (MICIU) and the Spanish Agencia Estatal de Investigación (AEI) MICIU/AEI/10.13039/501100011033 and co-financing from the European Regional Development Fund “ERDF A way of making Europe”. This work was also supported by grant This work was also supported by UNAM-PAPIIT IN221525 funded by the Universidad Nacional Autónoma de México (UNAM).

### Declaration of Competing Interest

The authors declare that they have no competing interests.

### CRediT authorship contribution statement

**Pieter van Dillewijn:** Conceptualization; Investigation; Data curation; Formal analysis; Visualization; Writing – original draft; Writing – review and editing. **Lydia M. Bernabéu-Roda:** Investigation; Writing – review and editing. **Virginia Cuéllar:** Investigation; Writing – review and editing. **Rafael Núñez:** Investigation; Formal analysis; Writing – review and editing. **Otto Geiger:** Writing – review and editing. **Isabel M. López-Lara:** Conceptualization; Writing – review and editing. **María J. Soto:** Conceptualization; Supervision; Funding acquisition; Writing – review and editing.

### Data availability

Data will be made available on request.

## Notes

### Competing Interest Statement

The authors have declared no competing interest.

