## Supplementary Material for "The effect of *Sinorhizobium meliloti* volatilomes and synthetic long-chain methylketones on soil and *Medicago truncatula* microbiomes"

**Section S1: Description of the microcosm experiments**

Microcosm experiments were set up using topsoil obtained from a site near to an experimental field of the Centro IFAPA Camino de Purchil in Granada in Southern Spain (37°10'20.7"N 3°38'11.5"W). Soil was sifted to 0.2 cm and stored until use at 4ºC. For experiments, sifted soil was mixed 1:1 (v/v) with sterile sand. The characteristics of this soil/sand mixture were the following: pH 8.55, assimilable potassium 70 p.p.m., assimilable phosphorus 8.2 p.p.m., organic nitrogen 0.04%, organic matter content 0.72%, and $CaCO_3$ content of 5.95%.

For soil experiments, a humidified soil/sand mixture was introduced into small pots (54 g/pot). The bottom of the pots was covered with a nylon mesh and set upon a small Petri dish (55 mm diameter) containing in the case of volatile methylketones (MKs) a Whatman filter paper (3MM) with 10 µl ethanol (control), 10 µl of a 50 mM solution of 2-TDC dissolved in ethanol, or of a mixture containing equal concentrations (50 mM each) of 2-TDC, 2-PDC and 2-HpDC dissolved in ethanol (MIX). In the case of volatilome experiments, the Petri dish contained solidified MM (1%) inoculated with 2 drops (10 µl each) of washed and concentrated cells of the *Sinorhizobium meliloti* GR4 (wt) cells, or its *fadA* mutant or with 2 drops of liquid MM (control). The Petri dishes were sealed under the pots with parafilm and introduced into a mini-greenhouse (Fig. S2a) with ventilation slots closed and sealed with masking tape. The mini-greenhouses were placed into a plant growth chamber with 22 ºC/ 18 ºC; 8 h day / 16 h night photoperiod and between 103-279 µmol $m^{-2}$ $s^{-1}$ lighting. Three replicate pots were taken initially (T=0) or after 1 and 7 days of incubation. To obtain soil samples, the contents of each pot were thoroughly mixed, placed in 50 mL tubes and stored at -20ºC until DNA extraction.

For plant assays, *Medicago truncatula* seeds were scarified, sterilized and germinated as described by Garcia et al. (2006). Small pots filled with 54 g/pot of the humidified soil/sand mixture were planted with germinated *M. truncatula* plantlets (1 plantlet/pot) and left to grow in a plant growth chamber under the conditions as above for 6 weeks. Fifteen plants of similar size and aspect were selected for either the volatile MKs or volatilome experiments. Three of the 15 selected plants were used as T=0 and 4 pots with underlying watering dish (Petri dish 55 mm diameter), were introduced into mini-greenhouses for each treatment (Fig. S2b). For volatile MKs, 4 small Petri dishes (55 mm diameter) each containing Whatman paper filters imbibed with 12.5 µl of 10% ethanol (control), 5 mM 2-TDC in 10% ethanol, or a mix with 5 mM of 2-TDC, 2-PDC and 2-

HpDC in 10% ethanol were introduced together with 4 planted pots. For volatilome experiments, a Petri dish (90 mm diameter) containing MM (1% agar) which had been plated either with 100 µl of washed and concentrated wt or *fadA* mutant cells or with 100 µl of liquid MM (control) was introduced into mini-greenhouses together with 4 planted pots (Fig. S2b). All mini-greenhouses were closed and sealed with masking tape and introduced into a plant growth chamber as described above. Plates with filters imbibed (or not) with MKs or plates inoculated (or not) were replaced after 1 week of incubation. After 14 days, plant pots were removed to obtain rhizosphere and root endosphere samples.

**Section S2: Obtainment of rhizosphere soil and root endosphere**

To obtain rhizosphere soil, roots of all the plants growing in the pots were gently shaken to remove excess bulk soil and then separated from the aerial part with sterile scissors before being introduced into a 15 ml or 50 ml tube containing 5 or 15 ml cooled sterile PBS buffer, respectively. Tubes were then vortexed for 1 min and introduced into a sonication bath (Bransonic ultrasonic cleaner) for 5 min. Roots were removed with sterile forceps to be used for endosphere determination and the tubes centrifuged at 6000 rpm on a table top centrifuge for 5 minutes. Supernatant was removed and the wet pellet considered to be rhizosphere soil was frozen at -20 °C for posterior DNA extraction. To obtain root endosphere samples, the roots were sterilized superficially by washing 1 minute with 70% ethanol followed by a wash with 3% sodium hypochlorite for 1 minute and three rinses with sterile distilled water. The roots were dried on sterile filter paper and frozen at -20ºC in an Eppendorf tube. Prior to DNA extraction, the sterilized roots were macerated into fine powder in liquid nitrogen.

**Section S3: Detailed Bioinformatic data analysis of 16S rRNA amplicon sequences**

16S rRNA amplicon sequences were processed using QIIME2 version 2024.5 (https://qiime2.org)) (Bolyen et al. 2019) in which forward and reverse sequences were trimmed, quality filtered, denoised, joined and dereplicated using DADA2 (Callahan et al. 2016). Feature frequency of original dataset was set to at least 10 repetitions. Representative features (a.k.a ASVs or amplified sequence variants) were classified with the SILVA database v138 (Quast et al. 2013), aligned with MAFFT (Katoh and Standley 2013) and phylogenetic trees were constructed using FastTree (Price et al. 2010). Features identified as mitochondria, chloroplasts or as Eukaryotes were removed. Alpha diversity indices (observed features and Shannon diversity index), and beta diversity of bacterial communities were determined using QIIME2. Principal coordinates analysis (PCoA) based on weighted Unifrac distances, as well as PERMANOVA analysis (999 permutations) and ADONIS analysis were also performed using QIIME2.

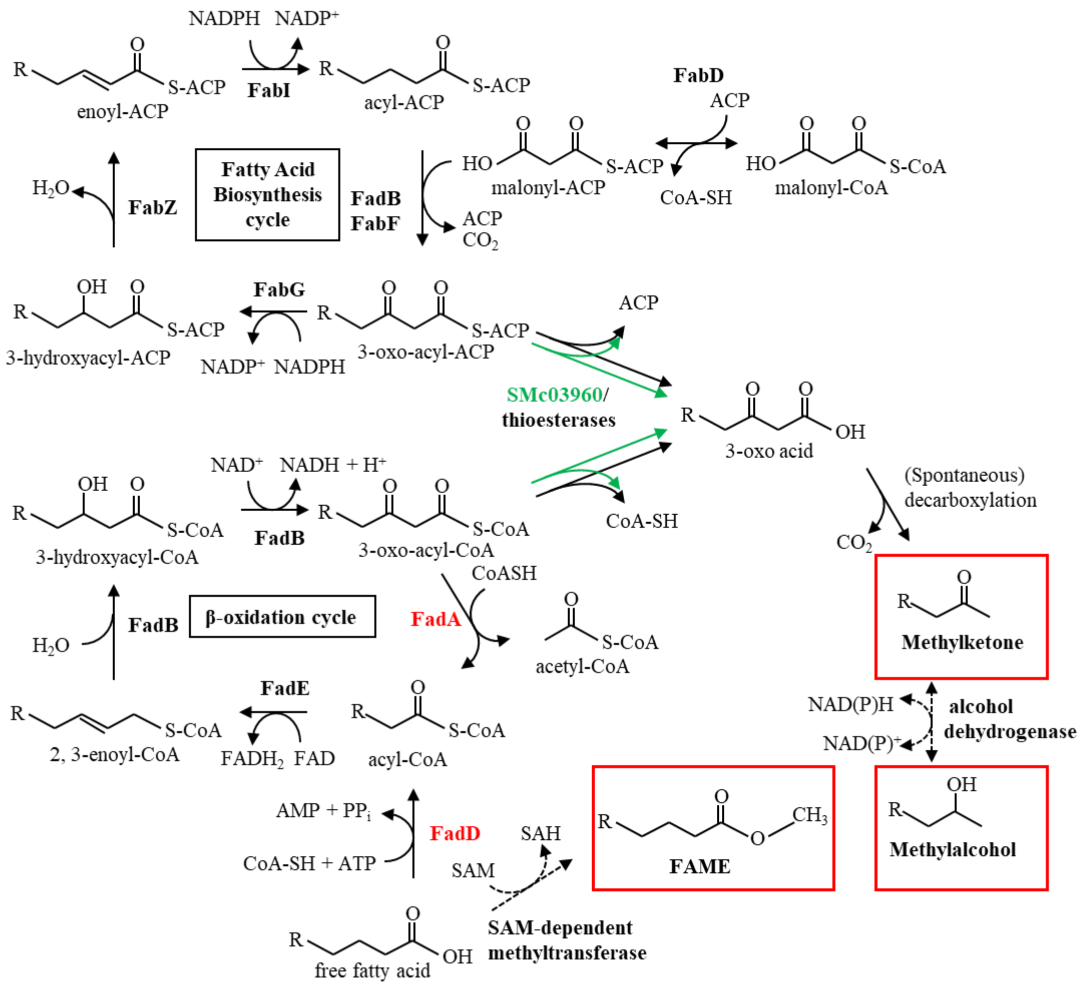


**Fig. S1** Fatty acid metabolism pathways related to the production of methylketones, methylalcohols and fatty acid methyl esters (FAMEs) (in red boxes). Enzymes targeted for disruption in mutants are indicated in red lettering. The elimination of FadD would lead to free fatty acid accumulation and disruption of FadA to the accumulation of 3-oxo-acyl-CoA. The reactions catalyzed by the thioesterase SMc03960 (Bernabéu-Roda et al., 2025) are indicated in green. Hypothetical reactions are indicated as dotted lines. ACP: acyl carrier protein; CoA: coenzyme A; SAM: *S*-adenosylmethionine; SAH: *S*-adenosylhomocysteine.

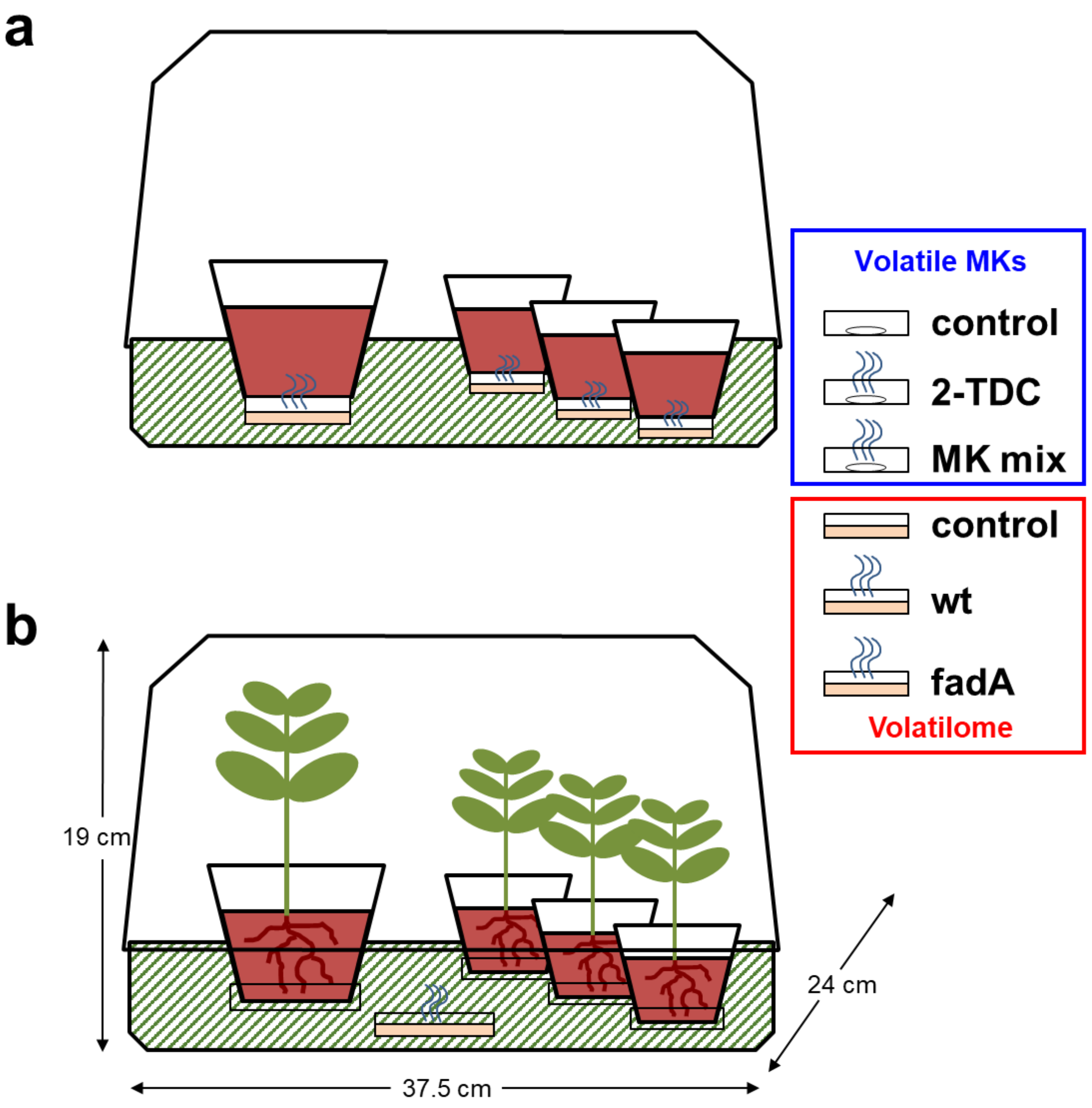


**Fig. S2** Experimental set-up in mini-greenhouses used to analyze the effect of Sm volatilomes and volatile methylketones (MKs) on the bacterial communities of a) soil and b) plants of *Medicago truncatula*. On the right, for the volatile MK experiments Petrie dishes contained filters (control) or impregnated with 2-tridecanone (2-TDC) or an MK mix of 2-TDC, 2-PDC and 2-HpDC. For the volatilome experiments Petrie dishes contained medium (control) or inoculated with GR4 (wt) or the *fadA* mutant (fadA).

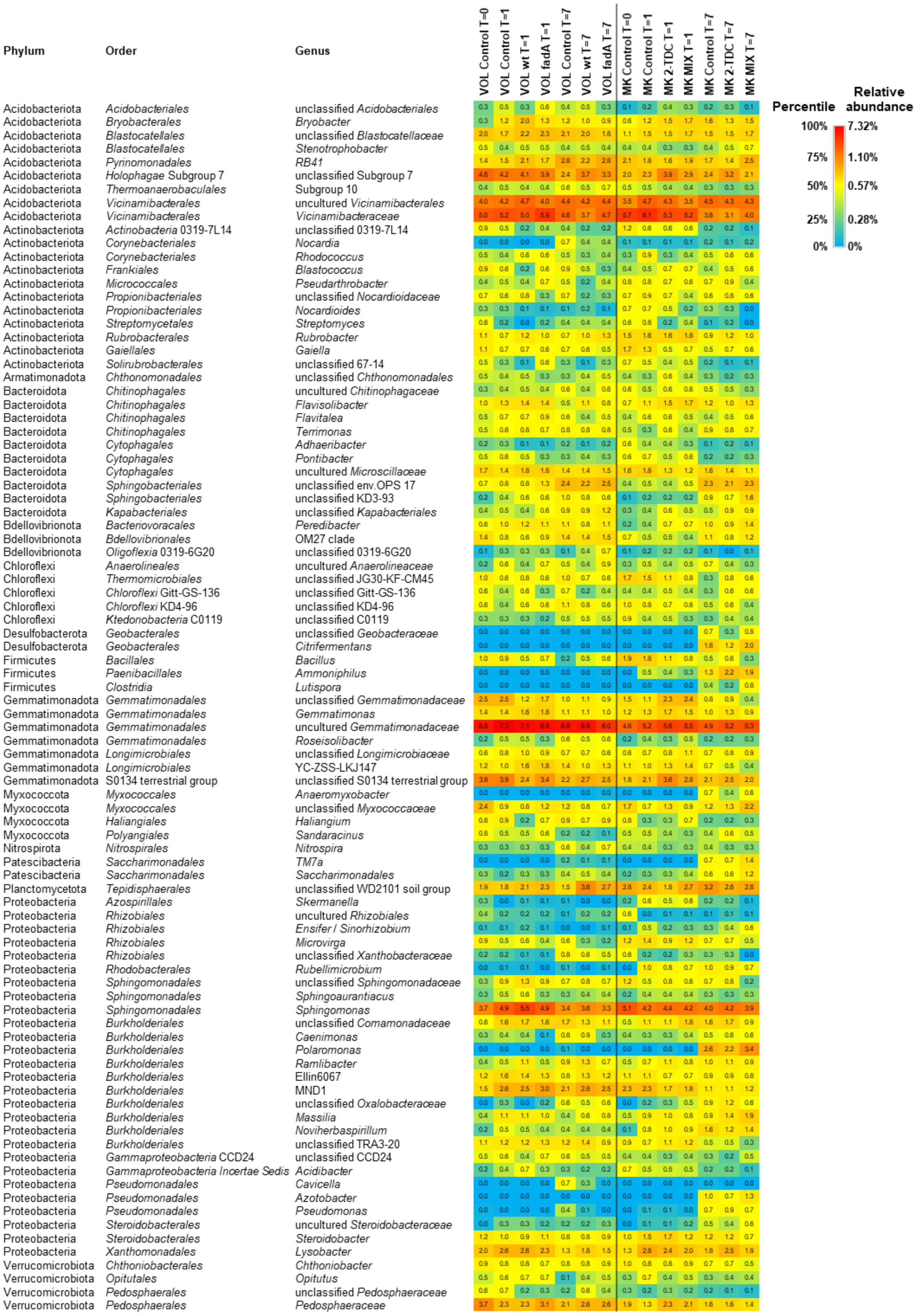


**Fig. S3** Heatmap of mean relative abundances of the 90 most abundant bacterial genera (≥0.57 % abundance in any condition) of replicate samples for each condition and time in soils exposed to Sm volatilomes (VOL) or to volatile methylketones (MK). MIX indicates the mixture of 2-TDC, 2-PDC and 2-HpDC.

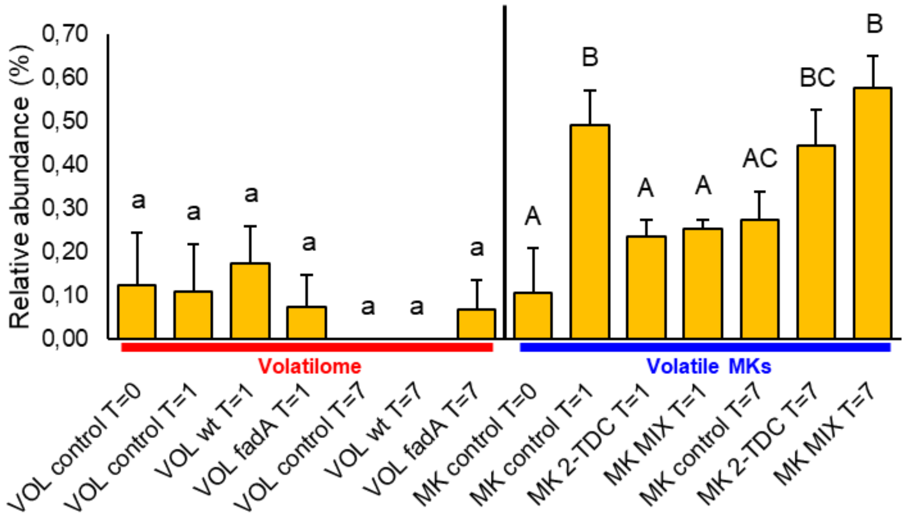


**Fig. S4** Relative abundance of *Ensifer*/*Sinorhizobium* in soils exposed to Sm volatilomes (VOL) or to volatile methylketones (MK). MIX indicates the mixture of 2-TDC, 2-PDC and 2-HpDC. Error bars indicate standard error and letters indicate significant differences (Welch T-test; $p<0.05$).

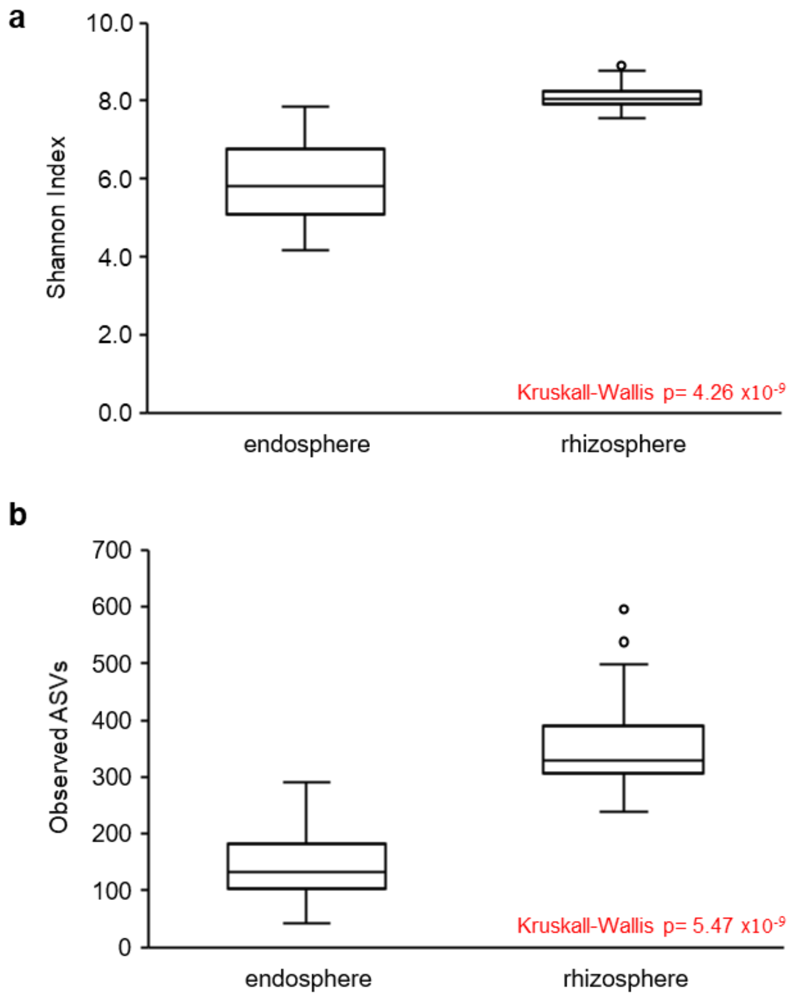


**Fig. S5** Box and whisker plots of alpha diversity indices for all root endosphere and rhizosphere samples. a) Shannon Index for bacterial diversity. b) Richness according to the number of observed ASVs. The middle line of the box and whisker plot represents the median. The bottom line of the box represents the median of the 1st quartile. The top line of the box represents the median of the 3rd quartile. The whiskers (vertical lines) extend from the ends of the box to the minimum and maximum values. Circles indicate outlier values.

| Phylum | Order | Genus | VOL control T=0 | VOL control T=14 | VOL wt T=14 | VOL fadA T=14 | MK control T=0 | MK control T=14 | MK 2-TDC T=14 | MK MIX T=14 |
|---|---|---|---|---|---|---|---|---|---|---|
| Acidobacteriota | *Bryobacterales* | *Bryobacter* | 0.7 | 0.7 | 0.3 | 0.7 | 0.3 | 0.6 | 1.1 | 0.7 |
| Acidobacteriota | *Blastocatellales* | unclassified *Blastocatellaceae* | 1.4 | 0.7 | 1.2 | 0.9 | 1.3 | 1.2 | 1.3 | 1.7 |
| Acidobacteriota | *Pyrinomonadales* | RB41 | 1.0 | 0.9 | 0.3 | 0.6 | 1.3 | 0.6 | 0.5 | 0.6 |
| Acidobacteriota | *Vicinamibacterales* | uncultured *Vicinamibacterales* | 3.8 | 3.6 | 3.0 | 2.0 | 2.9 | 1.8 | 2.9 | 2.6 |
| Acidobacteriota | *Vicinamibacterales* | unclassified *Vicinamibacteraceae* | 3.3 | 2.8 | 2.6 | 2.4 | 2.4 | 1.0 | 2.7 | 1.7 |
| Actinobacteriota | *Frankiales* | *Blastococcus* | 1.1 | 0.6 | 0.7 | 0.9 | 1.2 | 0.6 | 1.0 | 0.8 |
| Bacteroidota | *Chitinophagales* | unclassified *Chitinophagaceae* | 0.6 | 1.7 | 1.5 | 1.1 | 1.0 | 1.6 | 0.8 | 1.3 |
| Bacteroidota | *Chitinophagales* | *Terrimonas* | 1.4 | 1.7 | 1.2 | 0.8 | 0.9 | 1.5 | 1.3 | 1.4 |
| Bacteroidota | *Chitinophagales* | uncultured *Chitinophagaceae* | 0.8 | 1.9 | 1.7 | 0.6 | 0.8 | 1.3 | 0.5 | 1.2 |
| Bacteroidota | *Chitinophagales* | uncultured *Saprospiraceae* | 0.6 | 0.9 | 1.4 | 0.5 | 0.8 | 1.0 | 0.8 | 1.0 |
| Bacteroidota | *Cytophagales* | *Ohtaekwangia* | 1.5 | 1.5 | 2.2 | 1.6 | 1.2 | 1.6 | 1.7 | 1.0 |
| Bacteroidota | *Cytophagales* | uncultured *Microscillaceae* | 1.2 | 0.9 | 1.5 | 1.1 | 1.0 | 1.4 | 1.0 | 1.3 |
| Bacteroidota | *Flavobacteriales* | *Flavobacterium* | 1.1 | 1.6 | 1.9 | 2.1 | 1.0 | 2.0 | 0.8 | 1.0 |
| Bacteroidota | *Sphingobacteriales* | unclassified env.OPS_17 | 1.4 | 0.5 | 0.8 | 1.0 | 0.9 | 0.6 | 0.4 | 0.7 |
| Bacteroidota | *Kapabacteriales* | unclassified *Kapabacteriales* | 1.5 | 1.3 | 1.8 | 1.2 | 1.1 | 1.4 | 1.4 | 0.9 |
| Chloroflexi | *Chloroflexales* | uncultured *Roseiflexaceae* | 0.5 | 1.1 | 0.9 | 0.7 | 0.5 | 0.6 | 0.8 | 0.8 |
| Firmicutes | *Bacillales* | *Bacillus* | 1.1 | 0.5 | 0.6 | 0.6 | 0.9 | 0.5 | 0.6 | 1.0 |
| Firmicutes | *Clostridia* | unclassified *Gracilibacteraceae* | 0.2 | 1.2 | 0.3 | 0.7 | 0.5 | 1.4 | 1.0 | 0.5 |
| Firmicutes | *Clostridia* | *Lutispora* | 0.3 | 0.3 | 0.5 | 1.2 | 0.9 | 0.6 | 0.8 | 1.1 |
| Firmicutes | *Lachnospirales* | unclassified *Lachnospiraceae* | 0.2 | 0.9 | 0.6 | 0.9 | 0.6 | 1.2 | 0.6 | 0.9 |
| Firmicutes | *Desulfitobacteriales* | *Desulfosporosinus* | 0.4 | 1.2 | 0.1 | 1.0 | 0.4 | 0.6 | 1.3 | 1.0 |
| Firmicutes | *Negativicutes* | uncultured *Negativicutes* | 0.2 | 0.4 | 0.1 | 1.0 | 1.2 | 0.9 | 0.7 | 1.0 |
| Firmicutes | *Veillonellales-Selenomonadales* | *Pelosinus* | 0.6 | 1.4 | 0.7 | 1.5 | 0.7 | 1.1 | 1.7 | 1.1 |
| Firmicutes | *Veillonellales-Selenomonadales* | *Sporomusa* | 0.4 | 1.3 | 0.5 | 0.1 | 0.0 | 1.0 | 0.5 | 1.3 |
| Firmicutes | *Symbiobacteriales* | uncultured *Symbiobacteraceae* | 0.0 | 0.2 | 0.1 | 1.3 | 0.0 | 0.6 | 1.4 | 1.3 |
| Gemmatimonadota | *Gemmatimonadales* | uncultured *Gemmatimonadaceae* | 2.1 | 2.2 | 1.6 | 1.5 | 1.4 | 1.3 | 2.3 | 1.8 |
| Myxococcota | *Myxococcales* | unclassified *Myxococcaceae* | 1.0 | 0.4 | 0.5 | 0.8 | 0.7 | 0.5 | 0.3 | 0.4 |
| Myxococcota | *Haliangiales* | *Haliangium* | 1.0 | 0.3 | 0.3 | 0.5 | 1.4 | 0.8 | 1.2 | 1.1 |
| Planctomycetota | *Tepidisphaerales* | unclassified WD2101 soil group | 1.0 | 0.9 | 0.4 | 1.2 | 1.1 | 1.0 | 0.9 | 1.9 |
| Proteobacteria | *Rhizobiales* | *Devosia* | 0.2 | 0.7 | 1.3 | 0.6 | 0.7 | 1.3 | 1.1 | 0.8 |
| Proteobacteria | *Rhizobiales* | unclassified *Rhizobiaceae* | 0.8 | 1.4 | 1.8 | 1.6 | 0.7 | 1.1 | 0.6 | 0.8 |
| Proteobacteria | *Rhizobiales* | *Ensifer/Sinorhizobium* | 0.4 | 1.0 | 0.0 | 0.5 | 0.2 | 0.6 | 0.4 | 0.6 |
| Proteobacteria | *Rhodospirillales* | *Magnetospirillum* | 0.8 | 0.8 | 0.8 | 1.2 | 1.0 | 0.7 | 0.7 | 1.3 |
| Proteobacteria | *Sphingomonadales* | unclassified *Sphingomonadaceae* | 0.3 | 1.0 | 0.8 | 1.2 | 0.8 | 1.5 | 1.2 | 0.8 |
| Proteobacteria | *Sphingomonadales* | *Novosphingobium* | 0.3 | 0.6 | 1.0 | 0.8 | 0.6 | 1.4 | 0.6 | 0.7 |
| Proteobacteria | *Sphingomonadales* | *Sphingomonas* | 2.1 | 1.0 | 1.2 | 1.1 | 1.5 | 1.2 | 1.6 | 1.1 |
| Proteobacteria | *Burkholderiales* | unclassified *Comamonadaceae* | 1.6 | 1.3 | 2.2 | 2.3 | 0.9 | 1.8 | 1.6 | 1.6 |
| Proteobacteria | *Burkholderiales* | *Acidovorax* | 1.4 | 1.2 | 1.0 | 1.1 | 0.7 | 1.0 | 1.1 | 0.6 |
| Proteobacteria | *Burkholderiales* | *Rhizobacter* | 1.2 | 0.8 | 0.7 | 1.1 | 1.0 | 1.3 | 0.9 | 0.9 |
| Proteobacteria | *Burkholderiales* | *Methylotenera* | 0.8 | 0.7 | 1.1 | 0.5 | 0.7 | 1.2 | 1.0 | 0.9 |
| Proteobacteria | *Burkholderiales* | MND1 | 2.1 | 1.1 | 1.3 | 0.8 | 1.2 | 0.8 | 1.1 | 0.8 |
| Proteobacteria | *Burkholderiales* | *Massilia* | 1.2 | 0.8 | 0.5 | 1.0 | 1.5 | 0.9 | 1.4 | 1.0 |
| Proteobacteria | *Steroidobacterales* | *Steroidobacter* | 1.2 | 0.8 | 0.6 | 0.8 | 1.1 | 0.6 | 0.4 | 1.2 |
| Proteobacteria | *Xanthomonadales* | *Arenimonas* | 0.4 | 0.4 | 0.9 | 0.5 | 1.1 | 0.4 | 0.7 | 0.3 |
| Proteobacteria | *Pseudomonadales* | *Pseudomonas* | 0.2 | 0.1 | 0.0 | 0.2 | 0.2 | 0.2 | 0.1 | 0.0 |
| Verrucomicrobiota | *Opitutales* | IMCC26134 | 0.2 | 1.0 | 0.9 | 0.2 | 0.4 | 1.4 | 0.7 | 0.8 |
| Verrucomicrobiota | *Opitutales* | *Opitutus* | 0.3 | 1.3 | 1.2 | 1.1 | 0.8 | 1.6 | 1.5 | 1.0 |
| Verrucomicrobiota | *Pedosphaerales* | *Pedosphaeraceae* | 3.6 | 2.2 | 2.4 | 2.2 | 2.3 | 1.7 | 1.8 | 1.9 |
| Verrucomicrobiota | *Verrucomicrobiales* | *Luteolibacter* | 0.3 | 0.9 | 1.0 | 0.9 | 0.3 | 0.8 | 0.7 | 0.6 |

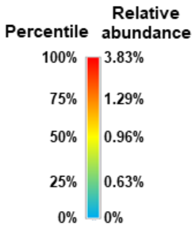


**Fig. S6** Heatmap of the mean relative abundances of the 49 most abundant bacterial genera (≥ 1% abundance in any condition) of replicate samples for each condition and time in the rhizosphere of plants exposed to Sm volatilomes (VOL) or to volatile methylketones (MK). MIX indicates the mixture of 2-TDC, 2-PDC and 2-HpDC.

| Phylum | Order | Genus | VOL control T=0 | VOL control T=14 | VOL wt T=14 | VOL fadA T=14 | MK control T=0 | MK control T=14 | MK 2-TDC T=14 | MK MIX T=14 |
|---|---|---|---|---|---|---|---|---|---|---|
| Actinobacteriota | *Corynebacteriales* | *Rhodococcus* | 0.0 | 1.0 | 0.6 | 0.7 | 0.0 | 0.6 | 0.3 | 0.5 |
| Actinobacteriota | *Micromonosporales* | unclassified *Micromonosporaceae* | 0.0 | 0.4 | 0.0 | 0.0 | 0.0 | 0.6 | 0.5 | 0.3 |
| Actinobacteriota | *Micromonosporales* | *Actinoplanes* | 0.5 | 0.4 | 0.6 | 0.1 | 0.2 | 0.0 | 0.2 | 0.5 |
| Actinobacteriota | *Streptomycetales* | *Streptomyces* | 1.2 | 2.8 | 1.0 | 0.8 | 1.0 | 2.7 | 1.8 | 2.2 |
| Bacteroidota | *Chitinophagales* | unclassified *Chitinophagaceae* | 0.5 | 1.9 | 1.1 | 1.0 | 0.7 | 1.3 | 1.3 | 1.3 |
| Bacteroidota | *Chitinophagales* | *Niastella* | 0.0 | 0.3 | 0.6 | 0.3 | 0.1 | 1.3 | 0.7 | 1.3 |
| Bacteroidota | *Chitinophagales* | uncultured *Chitinophagaceae* | 0.1 | 0.7 | 0.2 | 0.2 | 0.0 | 0.8 | 0.2 | 0.1 |
| Bacteroidota | *Cytophagales* | *Cytophaga* | 0.0 | 0.2 | 0.9 | 0.4 | 0.0 | 0.3 | 0.5 | 0.3 |
| Bacteroidota | *Cytophagales* | *Ohtaekwangia* | 0.1 | 0.5 | 0.6 | 0.6 | 0.5 | 2.3 | 2.2 | 1.6 |
| Bacteroidota | *Cytophagales* | *Dyadobacter* | 0.1 | 0.5 | 0.0 | 0.0 | 0.2 | 0.6 | 0.8 | 0.4 |
| Bacteroidota | *Flavobacteriales* | *Fluviicola* | 0.4 | 0.4 | 0.2 | 0.1 | 0.4 | 0.8 | 0.7 | 0.4 |
| Bacteroidota | *Flavobacteriales* | *Flavobacterium* | 0.4 | 2.2 | 0.8 | 0.5 | 1.8 | 3.2 | 3.9 | 2.7 |
| Bacteroidota | *Sphingobacteriales* | *Pedobacter* | 0.2 | 0.4 | 0.1 | 0.0 | 0.5 | 0.2 | 0.5 | 0.4 |
| Firmicutes | *Entomoplasmatales* | unclassified *Entomoplasmatales* type III | 0.0 | 0.0 | 0.5 | 0.6 | 0.0 | 0.1 | 0.2 | 0.0 |
| Firmicutes | *Paenibacillales* | *Paenibacillus* | 0.0 | 1.2 | 0.8 | 0.5 | 0.0 | 3.0 | 0.6 | 2.4 |
| Firmicutes | *Clostridia* | *Lutispora* | 0.0 | 0.7 | 0.1 | 0.2 | 0.1 | 0.6 | 0.2 | 0.9 |
| Firmicutes | *Clostridia* | unclassified *Hungateiclostridiaceae* | 0.0 | 0.6 | 0.6 | 0.0 | 0.0 | 0.4 | 0.6 | 0.4 |
| Firmicutes | *Clostridia* | *Anaerobacterium* | 0.0 | 0.3 | 0.1 | 0.3 | 0.0 | 0.7 | 0.3 | 1.2 |
| Firmicutes | *Clostridiales* | *Clostridium sensu stricto* 1 | 0.3 | 1.4 | 0.6 | 1.0 | 0.2 | 0.9 | 0.3 | 0.7 |
| Firmicutes | *Lachnospirales* | unclassified *Lachnospiraceae* | 0.2 | 1.7 | 0.6 | 0.4 | 0.0 | 2.4 | 0.4 | 1.4 |
| Firmicutes | *Lachnospirales* | uncultured *Lachnospiraceae* | 0.0 | 0.3 | 0.4 | 0.3 | 0.0 | 1.1 | 0.0 | 0.6 |
| Firmicutes | *Peptostreptococcales-Tissierellales* | *Anaerovorax* | 0.1 | 0.4 | 0.2 | 0.3 | 0.0 | 0.4 | 0.8 | 1.3 |
| Firmicutes | *Desulfitobacteriales* | *Desulfosporosinus* | 0.1 | 1.0 | 0.0 | 0.4 | 0.0 | 0.9 | 0.2 | 0.8 |
| Firmicutes | *Negativicutes* | uncultured *Negativicutes* | 0.0 | 0.8 | 0.1 | 0.8 | 0.3 | 1.5 | 2.3 | 1.7 |
| Firmicutes | *Veillonellales-Selenomonadales* | *Anaerosporomusa* | 0.1 | 0.6 | 0.0 | 0.0 | 0.0 | 0.6 | 1.0 | 1.0 |
| Firmicutes | *Veillonellales-Selenomonadales* | *Pelosinus* | 0.7 | 2.4 | 0.7 | 1.4 | 0.3 | 3.8 | 5.8 | 4.6 |
| Firmicutes | *Veillonellales-Selenomonadales* | *Sporomusa* | 0.1 | 1.7 | 0.5 | 0.2 | 0.0 | 2.1 | 0.8 | 2.2 |
| Planctomycetota | *Pirellulales* | Pir4 lineage | 0.1 | 0.2 | 0.3 | 0.2 | 0.1 | 0.3 | 0.5 | 0.9 |
| Proteobacteria | *Caulobacterales* | *Brevundimonas* | 0.0 | 0.4 | 0.0 | 0.0 | 0.2 | 0.8 | 0.6 | 0.5 |
| Proteobacteria | *Caulobacterales* | *Caulobacter* | 0.2 | 0.7 | 0.2 | 0.3 | 0.1 | 0.8 | 0.5 | 0.6 |
| Proteobacteria | *Rhizobiales* | *Devosia* | 0.0 | 0.5 | 0.4 | 0.3 | 0.2 | 1.6 | 1.1 | 1.1 |
| Proteobacteria | *Rhizobiales* | *Hyphomicrobium* | 0.0 | 0.2 | 0.5 | 0.0 | 0.0 | 1.1 | 0.5 | 1.2 |
| Proteobacteria | *Rhizobiales* | unclassified *Rhizobiaceae* | 0.2 | 1.6 | 0.8 | 0.5 | 0.3 | 1.1 | 1.8 | 2.0 |
| Proteobacteria | *Rhizobiales* | uncultured *Xanthobacteraceae* | 0.1 | 0.3 | 0.1 | 0.0 | 0.1 | 0.7 | 0.7 | 0.5 |
| Proteobacteria | *Rhodospirillales* | *Magnetospirillum* | 0.0 | 1.1 | 0.4 | 0.2 | 0.2 | 0.5 | 1.1 | 1.7 |
| Proteobacteria | *Sphingomonadales* | unclassified *Sphingomonadaceae* | 0.1 | 0.6 | 0.5 | 0.2 | 0.4 | 1.2 | 0.8 | 0.8 |
| Proteobacteria | *Sphingomonadales* | *Sphingomonas* | 0.4 | 0.4 | 0.7 | 0.2 | 0.4 | 1.0 | 0.2 | 0.9 |
| Proteobacteria | *Burkholderiales* | *Acidovorax* | 0.2 | 3.7 | 1.3 | 1.0 | 0.3 | 1.9 | 3.5 | 1.6 |
| Proteobacteria | *Burkholderiales* | *Azohydromonas* | 0.0 | 0.1 | 0.0 | 0.0 | 0.0 | 0.1 | 0.1 | 0.8 |
| Proteobacteria | *Burkholderiales* | *Pseudorhodoferax* | 0.2 | 0.2 | 0.0 | 0.0 | 0.3 | 0.8 | 0.4 | 0.6 |
| Proteobacteria | *Burkholderiales* | *Rhizobacter* | 0.0 | 0.8 | 0.9 | 0.7 | 0.1 | 0.3 | 0.5 | 0.2 |
| Proteobacteria | *Burkholderiales* | *Variovorax* | 0.1 | 0.5 | 0.8 | 0.1 | 0.3 | 0.3 | 0.4 | 0.0 |
| Proteobacteria | *Burkholderiales* | unclassified *Methylophilaceae* | 0.3 | 0.5 | 0.0 | 0.2 | 0.4 | 0.6 | 0.8 | 0.7 |
| Proteobacteria | *Burkholderiales* | *Methylotenera* | 0.4 | 0.5 | 0.2 | 0.3 | 0.3 | 0.3 | 0.7 | 0.5 |
| Proteobacteria | *Burkholderiales* | *Massilia* | 0.5 | 0.1 | 0.4 | 0.6 | 0.8 | 1.1 | 1.3 | 0.6 |
| Proteobacteria | *Enterobacterales* | unclassified *Enterobacteriaceae* | 0.0 | 0.4 | 0.0 | 0.0 | 0.0 | 2.2 | 0.0 | 0.3 |
| Proteobacteria | *Pseudomonadales* | *Acinetobacter* | 1.4 | 0.0 | 0.3 | 0.0 | 0.6 | 0.0 | 0.0 | 0.5 |
| Proteobacteria | *Pseudomonadales* | *Pseudomonas* | 0.1 | 1.0 | 1.6 | 0.7 | 0.5 | 1.1 | 1.4 | 0.4 |
| Proteobacteria | *Steroidobacterales* | *Steroidobacter* | 0.2 | 0.6 | 0.2 | 0.2 | 0.1 | 0.8 | 0.3 | 0.4 |
| Proteobacteria | *Xanthomonadales* | *Pseudoxanthomonas* | 0.2 | 0.5 | 0.0 | 0.0 | 0.1 | 0.6 | 0.3 | 0.0 |
| Verrucomicrobiota | *Opitutales* | IMCC26134 | 0.0 | 0.3 | 0.2 | 0.2 | 0.2 | 1.0 | 0.7 | 0.7 |
| Verrucomicrobiota | *Opitutales* | *Opitutus* | 0.4 | 1.2 | 0.2 | 0.5 | 0.7 | 1.1 | 0.8 | 1.6 |
| Verrucomicrobiota | *Verrucomicrobiales* | *Luteolibacter* | 0.0 | 0.5 | 0.1 | 0.0 | 0.2 | 0.6 | 0.3 | 0.7 |

| Percentile | Relative abundance |
|---|---|
| 100% | 5.83% |
| 75% | 0.77% |
| 50% | 0.41% |
| 25% | 0.15% |
| 0% | 0% |

**Fig. S7** Heatmap of the mean relative abundances of the 53 most abundant bacterial genera excluding *Ensifer*/*Sinorhizobium* ($\geq$ 0.5% abundance in any condition) of replicate samples for each condition and time in the root endosphere of plants exposed to Sm volatilomes (VOL) or to volatile methylketones (MK). MIX indicates the mixture of 2-TDC, 2-PDC and 2-HpDC.
